# Tear miRNAs identified in a murine model of Sjögren’s Syndrome as potential diagnostic biomarkers and indicators of disease mechanism

**DOI:** 10.1101/2021.10.10.463771

**Authors:** Shruti Singh Kakan, Maria C. Edman, Alexander Yao, Curtis T. Okamoto, Brooke E. Hjelm, Sarah F. Hamm-Alvarez

**Author notes:** Co-corresponding authors, Address correspondence to: Sarah F. Hamm-Alvarez, Department of Ophthalmology, 1450 San Pablo, Suite 4500, University of Southern California, Los Angeles CA 90033.

## Abstract

**Objective:** The tear miRNAome of the male NOD mouse, a model of ocular symptoms of Sjögren’s syndrome (SS), was analyzed to identify possible tear biomarkers of the disease.

**Methods:** Male NOD mice, aged 12-14 weeks, were used to identify tear miRNAs associated with development of autoimmune dacryoadenitis. Age-matched female NOD mice that do not develop the autoimmune dacryoadenitis characteristic of SS were used as negative controls while age- and sex-matched male BALB/c mice served as healthy controls. Total RNA was isolated from stimulated tears pooled from 5 mice per sample and tear miRNAs were sequenced and analyzed. Putative miRNA hits were validated using RT-qPCR in a separate mouse cohort, and the pathways influenced by the validated hits were identified using Ingenuity Pathway Analysis.

**Results:** In comparison to tears from both healthy (male BALB/c) and negative control (female NOD) mice, initial analysis identified 7 upregulated and 7 downregulated miRNAs in male NOD mouse tears. Of these, 8 were subsequently validated by RT-qPCR in tears from additional mouse cohorts. miRNAs previously implicated in SS pathology included mmu-miR-146a/b-5p, which were significantly downregulated in the male NOD mouse tears, as well as mmu-miR-150-5p and mmu-miR-181a-5p, which were upregulated in the male NOD mouse tears. All other validated hits including the upregulated miR-181b-5p and mmu-miR-203-3p, as well as the downregulated mmu-miR-322-5p and mmu-miR-503-5p, represent novel putative indicators of autoimmune dacryoadenitis in SS.

**Conclusions:** A panel of differentially expressed miRNAs were identified in tears of SS model male NOD mice, including some never previously linked to SS. These may have potential utility as diagnostic biomarkers for ocular symptoms of SS; evaluation of the pathways influenced by these dysregulated miRNAs may also provide further insights into SS pathogenesis.

## INTRODUCTION

Sjögren’s Syndrome (SS) is a chronic, progressive autoimmune disease that affects ~1% of the population(1) and causes inflammation of moisture-producing glands including lacrimal glands (LG) and salivary glands (SG), leading to dry eye and dry mouth(2). SS also causes systemic disease including inflammation in skin, lung, kidneys, and the nervous system(3) resulting in dryness of the skin, nose and vagina, debilitating muscle and joint pain, fatigue, and chronic cough(4). Among all autoimmune diseases, SS patients have the highest incidence of malignant lymphoma(5, 6). Diagnosis of SS is challenging because the current diagnostic criteria are cumbersome, involving several subjective and analytical tests requiring a blood draw and a painful and sometimes error-prone minor SG biopsy(7, 8). Additionally, symptoms of SS overlap with those of other autoimmune diseases such as rheumatoid arthritis (RA) and Systemic Lupus Erythematosus (SLE) and other dry eye diseases, leading to delays in diagnosis or misdiagnosis(9). Thus, it can take several years before the disease is confirmed, during which time infiltrating immune cells may further damage exocrine glands and sustain debilitating ocular and oral cavity symptoms. SS can also lead to other ocular complications such as uveitis and optic neuritis(10, 11). Although it is the second most common autoimmune disorder in the United States(12), SS receives far less attention for research and therapeutic development than does RA or SLE. With growing prevalence there is an urgent need for more specific diagnostic tests for SS to prevent irreversible tissue and/or organ damage and to improve the health and vision-related quality of life for SS patients.

An ideal diagnostic tear biomarker would be chemically stable and able to: 1) detect SS with high sensitivity and specificity; 2) distinguish SS from other autoimmune and dry eye diseases; 3) be collected relatively non-invasively; and 4) be processed inexpensively in a straightforward manner. With these characteristics in mind, we investigated a type of short noncoding RNAs called microRNA (miRNAs) that are 18-26 nucleotides long. miRNAs circulate in the body either packaged inside secreted vesicles or bound to RNA-binding proteins and are therefore highly stable and protected from RNAse degradation. miRNAs are responsible for transcriptional regulation of nearly 60% of all mammalian messenger RNA (mRNAs)(13), and dysregulation of miRNA has been implicated in diseases such as cancer and neurodegenerative disease(14, 15). Moreover, miRNAs and their mRNA targets have co-evolved and are highly evolutionarily conserved in mammals(16), allowing for studies in model organisms to be directly extrapolated to humans. This feature is critical for SS research, as mammalian model organisms allow investigation of the earliest stages of glandular inflammation, which is difficult to do in patients due to the substantial delay in SS diagnosis

While SS causes systemic symptoms, its hallmark manifestations are inflammation of LG and SG. Although other biofluids and their components, i.e., plasma/serum, peripheral blood mononuclear cells (PBMC) and saliva, have been extensively investigated as sources of protein and RNA based biomarkers, there are few reports of microarray-based assessment of tears for SS and none using comprehensive Next Generation Sequencing (NGS) analysis. Tears are highly enriched in miRNA compared to other biofluids such as serum and cerebrospinal fluid(17). Moreover, tear miRNA candidates have been investigated for detection of primary open-angle glaucoma(18) and Alzheimer’s disease(15), with very high sensitivity and specificity. Tear collection is quick, atraumatic, and non-invasive. Most importantly, changes in tear composition are likely to be directly related to the health of the LG, the principal source of most of the aqueous tear components and a primary target of autoimmune exocrinopathy(19). Thus, we propose that dysregulated tear microRNA may be indicative of LG disease in Sjögren’s syndrome and might also provide insights into disease pathogenesis.

This discovery study utilized male NOD mice, a well-established model of the ocular symptoms of SS. These mice exhibit lymphocytic infiltration of the LG, beginning around eight weeks of age. By 12 weeks of age, male NOD mice exhibit a notable lymphocytic infiltration of LG associated with decreased tear production, ocular surface inflammation, and increased secretion of cysteine proteases such as cathepsin S in tears(20, 21). In contrast, female NOD mice do not manifest marked ocular symptoms of SS, instead exhibiting SG inflammation at a later age of 16 weeks. They are thus typically used to study autoimmune sialadenitis. Age-matched female NOD mice were used as a negative control in our study because they lack significant LG disease and the minimal local LG disease that they may express occurs well beyond 30 weeks of age(22). As well, female NOD mice share the genetic background of the male NOD mice. Age- and sex-matched male BALB/C mice, which also lack LG disease and are frequently used as a control for male NOD and their derivatives in studying disease development(20, 23), served as a healthy control.

In this study, stimulated tears were collected from each mouse cohort and pooled for isolation of total RNA, prior to conducting a comprehensive small RNA sequencing analysis (**Supplemental Fig 1**). Over 500 distinct miRNAs were identified in male NOD mouse tears of which 14 were differentially expressed compared to tears from both control groups. These findings were validated in separate cohorts of mice using qRT-PCR, confirming differential expression of 8 miRNAs. In tears of 12-14 weeks old male NOD mice, miR-146a-5p, miR-146b-5p, miR-322-3p, and miR-503-5p were down-regulated; whereas miR-181a-5p, miR-181b-5p, miR-150-5p, and miR-203-3p were up-regulated. To our knowledge, this is the first comprehensive tear miRNA sequencing analysis in an animal model of SS, the results of which may have important implications for clinical diagnosis of ocular involvement in SS patients.

## MATERIALS & METHODS

### Animals

Male NOD/ShiLtJ (001976) mice aged 12-14 weeks were used as an early-intermediate disease model for ocular manifestations of SS(20, 24). While age-matched male BALB/cJ (000665) mice served as a healthy control, female NOD/ShiLtJ mice that <u>do not</u> develop lymphocytic infiltration in the LG or other ocular manifestations of Sjögren’s syndrome by 12 weeks were employed as an additional negative, strain-specific control(22). Mice older than 14 weeks were excluded from the study to avoid potential confounding effects of diabetes. All strains were purchased from Jackson Laboratories (Bar Harbor, Maine). All animal procedures and experiments followed protocols approved by the Institutional Animal Care and Use Committee (IACUC) at the University of Southern California (Protocol #10547) as well as the Guide for Care and Use of Laboratory animals, Eighth Edition(25). All sections of this report adhere to the ARRIVE Guidelines for reporting animal research and a completed ARRIVE guidelines checklist is included in Checklist S1.

### Tear and tissue collection

Stimulated tears were collected as described previously(20). Briefly, mice were anesthetized with 20 mg/kg of Ketamine and 2 mg/kg of Xylazine and placed on heating pads. An incision was made in the skin above the LG, and the exposed gland was stimulated with 3 μL of 50 μM carbachol (Alfa Aesar, Haverhill, MA). Tears were collected using 2 μL sized Microcaps glass capillaries (Drummond Scientific, Broomall, PA) for 5 min after topical stimulation. This process was repeated twice for a total of 15 min of stimulated collection, after which the mice, still under anesthesia were euthanized by cervical dislocation. One LG (either left or right) was selected at random for RNA isolation and the remaining LG was processed for histology using Hematoxylin and Eosin (H & E) staining. Tears were pooled from 5 mice for a single sample intended for Next Generation Sequencing (NGS) (to allow collection of sufficient RNA amounts for NGS library preparation), with 5 samples each for male NOD and BALB/c mice and 3 samples for female NOD mice. For RT-qPCR validation experiments, tears were similarly collected; however, tears comprising one sample for this validation were pooled from 3 mice from each mouse group. Four pooled samples per group were analyzed. From each mouse, one LG was collected at random for RNA isolation and the other for histology. Cost constraints are the primary reason for the choice of the sample size for the discovery and validation cohorts. Number of mice used per sample to pool tear RNA was determined by the minimum amount of RNA required for NGS library preparation (>10 ng RNA per sample) or RT-qPCR (> 1 ng RNA per sample).

### LG Histology & Quantification of Lymphocytic Infiltration

LGs were placed in cassettes which were then fixed in 10% neutral buffered formalin solution for at least 24 h and transferred to 70% ethanol. Tissues were embedded in paraffin, and longitudinal cross-sections of 5-6 μm thickness were cut at various depths into the LG. Each tissue section was stained with H&E dye and images were acquired by a AxioScope 5 with Axiocam 305 microscope (Carl Zeiss AG, Jena, Germany). Infiltrates were quantified using Image J(26) to calculate the total area of the LG and the area occupied by infiltrating lymphocytes. The percentage infiltration was calculated by dividing the area occupied by foci divided by the total area of the cross-section multiplied by 100. The percentage of infiltration at three depths within the LG in each mouse was averaged to reflect the total amount of lymphocytic infiltration/gland.

### RNA Isolation, Quality Control, Library Preparation and NGS

5 μL of β-mercaptoethanol (BME) was added to each pooled mouse tear sample to prevent degradation by RNAase. Addition of BME to tears improved RNA quality as indicated by improvement of Agilent Tapestation traces and RNA integrity Number (RIN) (data not shown). Total RNA was isolated using the miRNEasy total RNA isolation kit (Qiagen) following the manufacturer’s guide, with a slight change in the final step. RNA was eluted from the dried column by adding 14 μL of RNAse free water and eluting twice for a total elution volume of ~28 μL. RNA was stored at −80°C prior to library preparation. RNA quality and concentration was assessed in house using Tapestation with a High Sensitivity RNA ScreenTape (5067-5579, Agilent) to ensure the RNA was of high integrity and sufficient quantity for NGS library preparation. Samples with RIN scores lower than 2 or RNA < 10 ng were excluded from sequencing.

Library preparation and NGS was outsourced to Qiagen where the RNA quality was again assessed using Nanodrop and Tapestation prior to library preparation. Order of samples for library preparation, and NGS was not known to the authors. The small RNA library was prepared using the QIAseq miRNA library kit (331502, Qiagen), specifically optimized for very low input small RNA, following the manufacturer’s guidelines. Small RNA sequencing was conducted on an Illumina Nextseq 500/550 high output flow cell with 75 cycles in single end configuration with an average sequencing depth of 17 to 18 million reads/sample (expected to be sufficient to provide >1 million mapped miRNA reads per sample to allow sufficient power to detect differentially expressed miRNA in accordance with exRNA consortium guidelines)

### Bioinformatics analysis

Read quality of the FASTQ files from all samples were assessed using FASTQC; adapter and quality trimming was done with Cutadapt v2.0. Trimmed reads were aligned to mouse whole genome GENCODE GRCm38 using Bowtie and quantified using featureCounts. Raw reads were also aligned to miRbase v 22.0 as described previously(27) using our in-house aligner miRGrep(27). miRNA read counts filtering, data normalization, and differential gene expression analysis was done using DESEq2 in RStudio with no data points excluded in the analysis. (A full list of software packages, R Code and program parameters can be found in **Supplemental Fig S1**).

### RT-qPCR Validation

Validation of miRNA hits from the tear miRNA library utilized RT-qPCR analysis of tear miRNA isolated from separate cohorts of mice using Qiagen’s Custom PCR Panels. 13 miRNAs were tested. Due to constraints in designing the custom PCR panel to validate the original 14 hits, miR-322-3p was not included. We reasoned that if the 5p counterpart, evaluated in the panel, was confirmed as differentially expressed, it would be reasonable to expect that the 3p counterpart might also be differentially expressed since they are the products of the same precursor miRNA, miR-322. Real time PCR was carried out in triplicate on a QuantStudio 6 Flex instrument (Applied Biosystems, Foster City, CA, USA). Raw Ct values were first normalized using the inter-plate calibrator UniSp3, followed by normalization to internal controls, miR-25-3p and miR-16-5p. Statistics were performed on ΔCt values. Results were reported with respect to both negative and healthy controls using the ΔΔCt method employing QuantStudio Real-Time PCR system software v. 1.3.

### Ingenuity Pathway analysis of miRNA hits

miRNA hits validated by qPCR were uploaded to Ingenuity Pathway Analysis (IPA) with fold-change and p-values calculated from DESeq2. Using the ‘microRNA target filter’ module of IPA, a list of mRNA targets sourced from TargetScan was obtained. IPA recognized seven microRNA hits and reported a list of 3991 mRNA targets. This list was shortlisted by a) confidence (experimentally validated, predicted high) to 1100 mRNA targets, and b)species (restricted to mammals only–human, mouse, and rat), from which, a network map was generated. Using the build and connect features of the main ‘pathways’ module, mRNA targets involved in immunological and inflammatory diseases, inflammatory responses, and cancer were shortlisted. Pathways, disease categories and functions over-represented by the miRNA and their mRNA targets were obtained using the overlay feature of the ‘pathways’ module in decreasing order of probability. From here, pathways most significantly over-represented were evaluated to gain further insights into the potential role of the dysregulated miRNA hits in SS.

## RESULTS

### Histological Analysis of LG lymphocytic infiltration

To determine the percentage of lymphocytic infiltration in mouse LG, H&E-stained images of LG sections from the mice used for the initial tear collection were acquired and analyzed. Neither the LG of female NOD mice (**Figure 1A**) or healthy male BALB/C mice (**Figure 1B**) at 12 weeks of age showed any lymphocytic infiltration, as expected(22). The male NOD LG had notable lymphocytic infiltration by 12 weeks of age (**Figure 1C**), confirming establishment of LG disease associated with SS. The percentage infiltration of LG from the male NOD mice was thus significantly higher than both the male BALB/c and female NOD mice in both discovery (**Figure 1D**) and validation (**Figure 1E**) cohorts.

**Figure 1.**
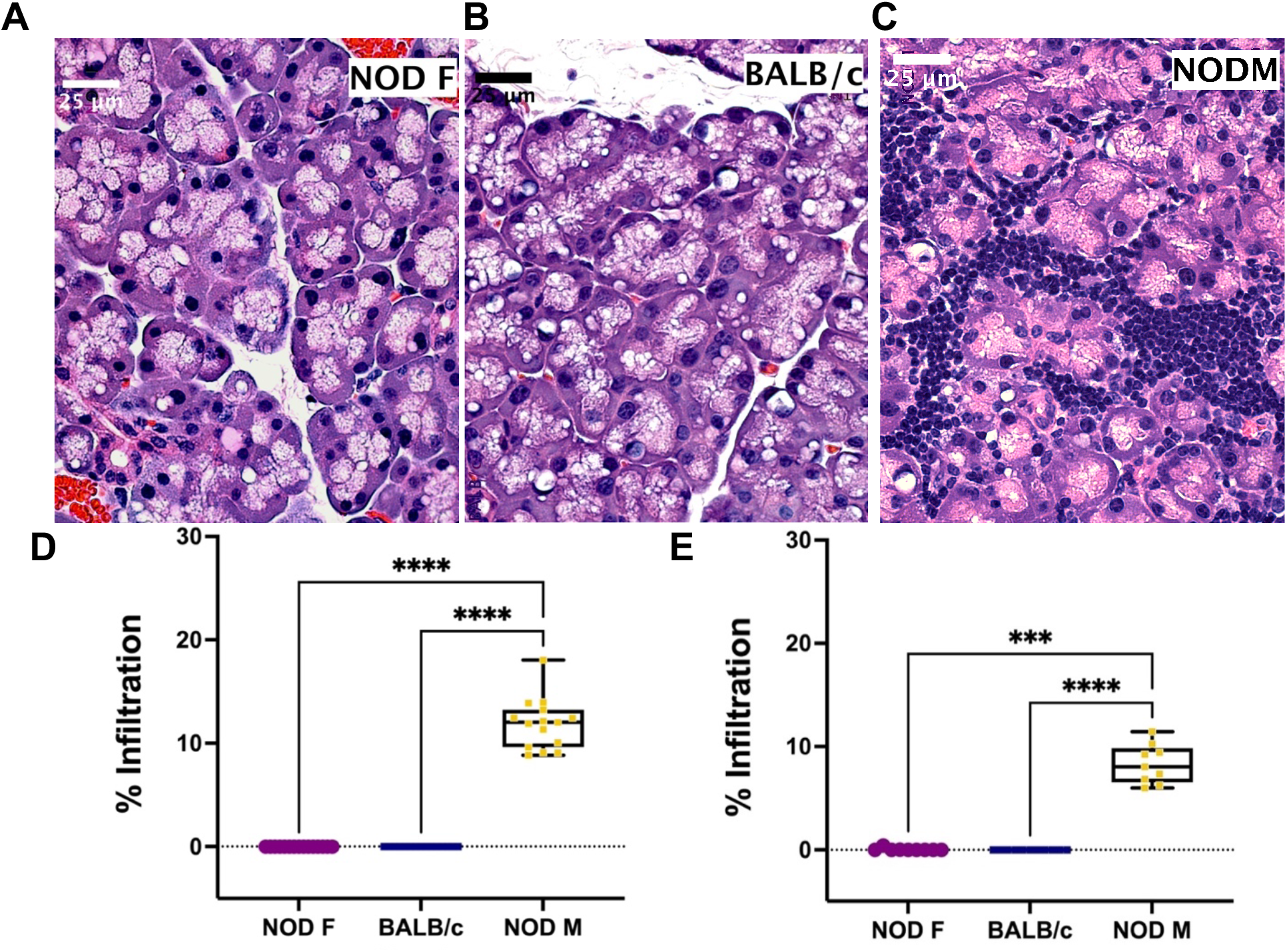
Representative H & E-stained images of LG from 12-14-week-old female NOD, male BALB/c or male NOD mice. No lymphocytic infiltration is observed in LG from (A) female NOD mice or B) male BALB/c mice. Marked lymphocytic infiltration is observed in LG from (C) male NOD mice. Scalebar, 25 μm. The percentage of total area of infiltrating lymphocytes in LG calculated in both the (D) discovery cohort and (E) validation cohort of age-matched female NOD mice (purple), male BALB/c mice (blue) and male NOD mice (yellow). Values for the average percentage lymphocytic infiltration per one LG per mouse are plotted as a boxplot with whiskers showing range. N=5 samples for male NOD and BALB/c, and N=3 samples for female NOD mice; n=5 mice per sample (for the discovery cohort), and n=3 mice per sample (for the validation cohort). Points represent data for individual mouse glands. (*** p < 10^−3^, **** p < 10^−4^, Kruskal-Wallis ANOVA).

### Male NOD mice have differential tear production, but not total protein or RNA yields

As the LG is the predominant source of aqueous tear components, changes in miRNA composition of tears may be indicative of LG disease. We have established previously that male NOD mice have decreased tear production, similarly to SS patients(20, 28). However, their tear protein concentration when normalized to tear volume is not affected(20). Similarly, we did not observe a significant difference in the RNA tear concentration normalized to tear volume in the male NOD mice as compared to male BALB/c and female NOD mice (**Supplemental Fig S2A**). There were also no significant differences in the RNA concentration or quality, as indicated by the RNA integrity number (RIN) of samples in each group (**Supplemental Fig S2B**). Assessing the total number of reads aligned to miRbase v22, the counts per million miRNA or CPM miRNA normalized to RNA amount per unit tear volume, showed no significant difference across the three groups (**Supplemental Fig S2C**).

### Male NOD mice tears from the discovery cohort exhibit differentially expressed miRNA

We detected 563 distinct miRNAs in male NOD mouse tears, 511 miRNAs in the BALB/c mouse tears, and 622 miRNAs in the female NOD mouse tears (**Figure 3A**). About 455 tear miRNAs were common to all three strains while 28 and 21 miRNAs were unique to male NOD and male BALB/c mouse strains, respectively. Female NOD mouse tears had the greatest number of distinct miRNAs at 84, which is likely a sex effect. Female NODs also had more tear miRNA in common with male NOD (64) than with male BALB/c (19), which may be attributed to a strain effect. Both sex and strain were used as co-variates in the downstream statistical analysis.

A miRNA was considered as a ‘hit’ if it had 1) a normalized base mean expression of at least 10 reads, 2) was up or downregulated in male NOD tears by at least ± 0.5 Log2 Fold Change (L2FC) in the same direction when compared to both control tear samples from male BALB/c and female NOD tears, and 3) resulted in a significant (p<0.05), unadjusted p-value in at least one of the two comparisons. With these criteria, using DESEq2 we identified 14 miRNAs as differentially expressed (**Table 1**). Seven miRNAs (miR-181a-5p, miR-181b-5p miR-203-3p, miR-150-5p, miR-3076-3p, miR-3963, miR-3572-3p) were upregulated (**Figure 2A**), while another 7 miRNAs (miR-146a/b-5p, miR-147-3p, miR-322-3p, miR-322-5p, miR-421-3p, miR-503-5p) were downregulated in tears of male NOD mice (**Figure 2B**).

**Figure 2.**
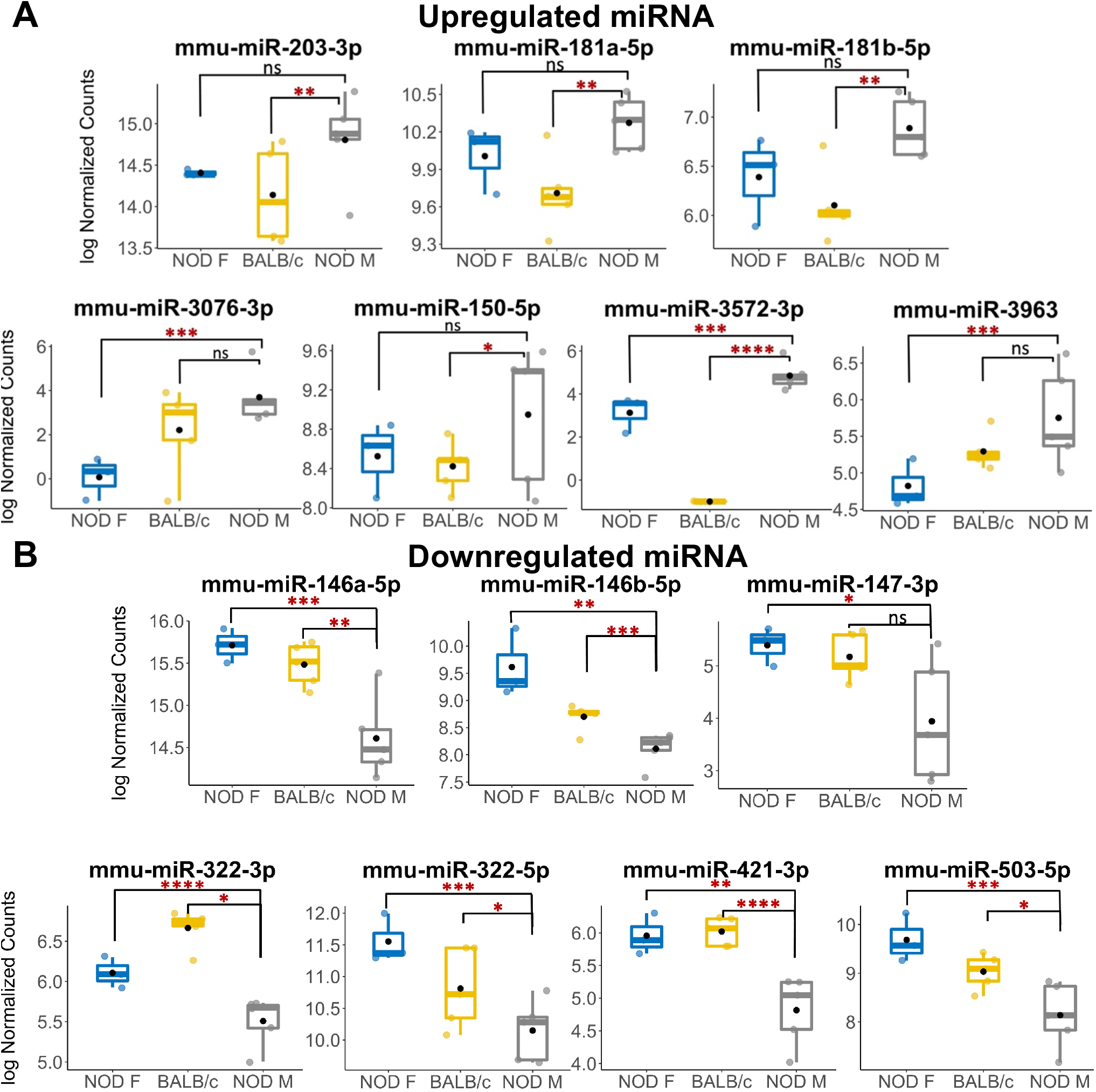
Multiple miRNAs are differentially expressed in tears of diseased male NOD mice. (A) 7 miRNAs were identified as upregulated in tears of 12 week male NOD mice while (B) 7 more miRNAs were found to be downregulated in tears of 12-14 week male NOD mice as compared to both age-matched female NOD and male BALB/c mice. Log10 Normalized miRNA counts as calculated by DESeq2 are plotted for each strain. N=5 samples for male NOD and BALB/c, and N=3 samples for female NOD mice; n=5 mice per sample for each strain. miRNA were considered differentially expressed if the fold change trended in the same direction for NOD M vs BALB/c and NOD M vs NOD F; had a mean expression value of 10 reads or higher and had a significant unadjusted p value in at least one of the two comparisons. (* p < 0.05; ** p < 0.01, *** p < 0.001, **** p < 0.0001, DESeq2).

**Table 1.** Differentially expressed miRNA in male NOD mice tears as compared to those in tears of male BALB/c and female NOD mice.

| miRNA | Log2 Fold Change |  | P<sub>value</sub> |  |
| --- | --- | --- | --- | --- |
|  | NOD M vs. NOD F | NOD M vs. BALB/c M | NOD M vs. NOD F | NOD M vs. BALB/c M |
| mmu-miR-203-3p | 0.656 | 0.477 | **0.0389** | 0.1937 |
| mmu-miR-181b-5p | 0.779 | 0.432 | **$2.964 \times 10^{-3}$** | 0.1569 |
| mmu-miR-181a-5p | 0.547 | 0.258 | **$6.075 \times 10^{-3}$** | 0.2634 |
| mmu-miR-3076-3p | 1.269 | 4.143 | 0.0959 | **$3.810 \times 10^{-4}$** |
| mmu-miR-150-5p | 0.639 | 0.510 | **0.0473** | 0.1713 |
| mmu-miR-3572-3p | 7.245 | 1.583 | **$1.270 \times 10^{-12}$** | **$9.008 \times 10^{-3}$** |
| mmu-miR-3963 | 0.595 | 0.999 | 0.0903 | **0.0182** |
| mmu-miR-146a-5p | -0.826 | -1.045 | **$1.141 \times 10^{-3}$** | **$3.631 \times 10^{-4}$** |
| mmu-miR-146b-5p | -0.578 | -1.560 | **0.02774** | **$2.406 \times 10^{-7}$** |
| mmu-miR-147-3p | -0.967 | -1.178 | 0.0541 | **0.0427** |
| mmu-miR-322-3p | -0.544 | -1.136 | **0.0423** | **$3.526 \times 10^{-7}$** |
| mmu-miR-322-5p | -1.376 | -0.707 | **$4.558 \times 10^{-4}$** | **$3.764 \times 10^{-3}$** |
| mmu-miR-421-3p | -1.125 | -1.028 | **$4.470 \times 10^{-5}$** | **$1.450 \times 10^{-3}$** |
| mmu-miR-503-5p | -0.812 | -1.485 | **0.0165** | **$1.459 \times 10^{-4}$** |

*p-values and log2FoldChanges were calculated using DESeq2. Significant unadjusted p-values are in bold.*

Unsupervised hierarchical clustering analysis (shown as heatmaps) of differentially expressed miRNA clustered female NOD with BALB/c separately from male NODs (**Figure 3B**). Sample level variation is illustrated in the heatmaps. Principal component analysis (PCA) of the data shows considerable overlap between male NOD and BALB/c when comparing all miRNAs, while the female NOD samples cluster away from the males. PC1 accounts for 41% of the variation in the data, likely attributable to sex (**Figure 3C**). When comparing the top miRNA hits, however (**Figure 3D**), there do not appear to be any outliers and samples within a group correlate well. PC1 accounts for 58% of the variation and appears to show a strain effect with controls clustering close to each other and away from male NODs. PC2 accounts for 23% of the variation and shows a sex effect. Volcano plots comparing tear miRNA expression show greater variability between male and female NOD mice (**Figure 3F**) and less so in male NOD and BALB/c mice tears (**Figure 3E**).

**Figure 3.**
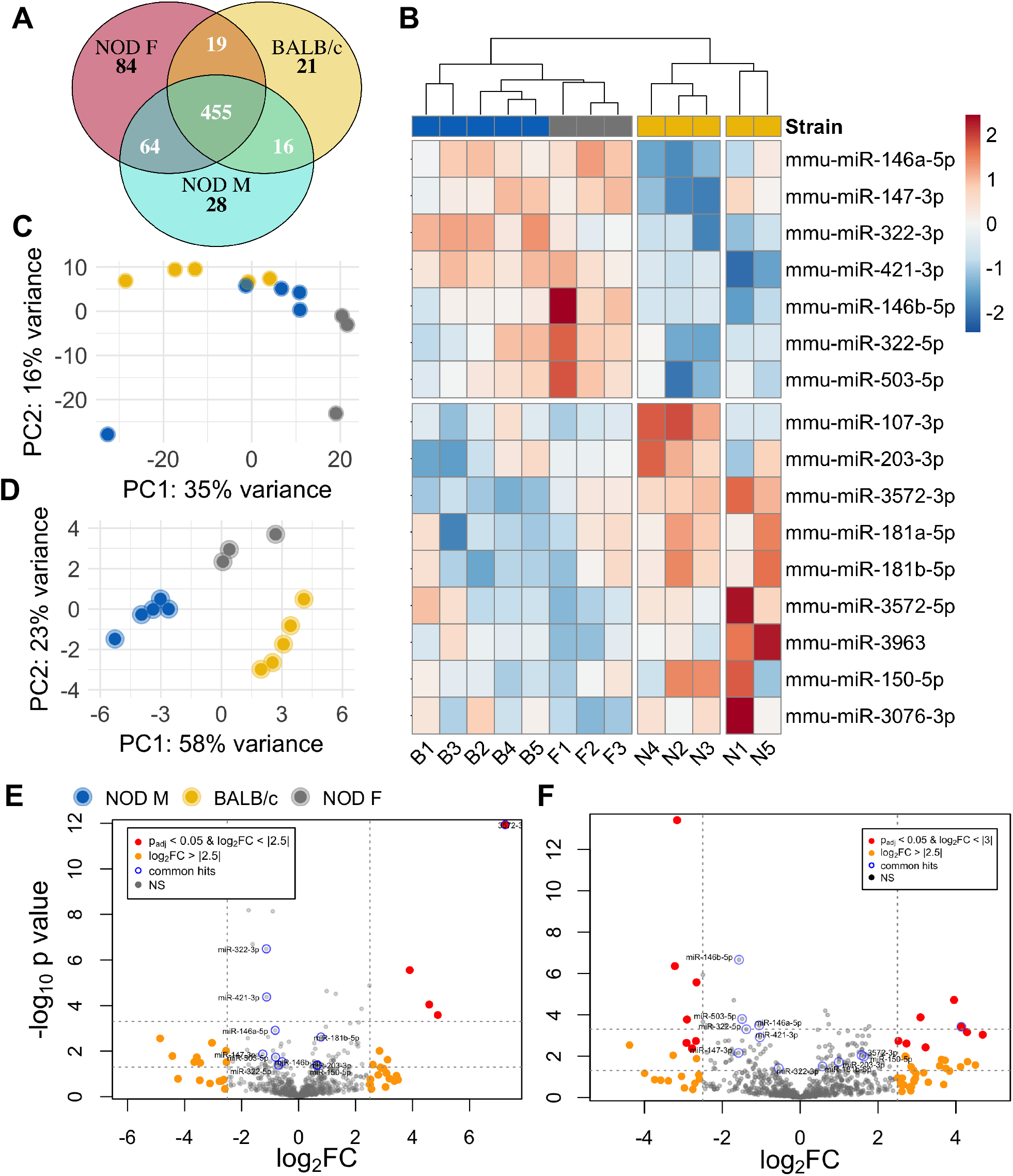
Differential miRNA expression analysis of NOD mouse tears. (A) Venn diagram showing the overlap of miRNA in the three strains. miRNA that had normalized read counts > 0 in at least 4 out of 5 samples of NOD and BALB/c and 2 out of 3 samples for the female NOD were included for calculations. (B) Heatmap of miRNA that are differentially expressed in male NOD mouse tears as compared to tears of male BALB/c and female NOD. Principal component analysis (PCA) plot of the (C) complete miRNA data and (D) top miRNA hits, characterizes the trends exhibited by the expression profiles of the 3 strains. Each dot represents a sample, and each color represents the type of the sample. Volcano plot of differentially expressed miRNA in NOD mouse tears as compared to (E) that of male BALB/c and (F) female NOD.

**Figure 3.**
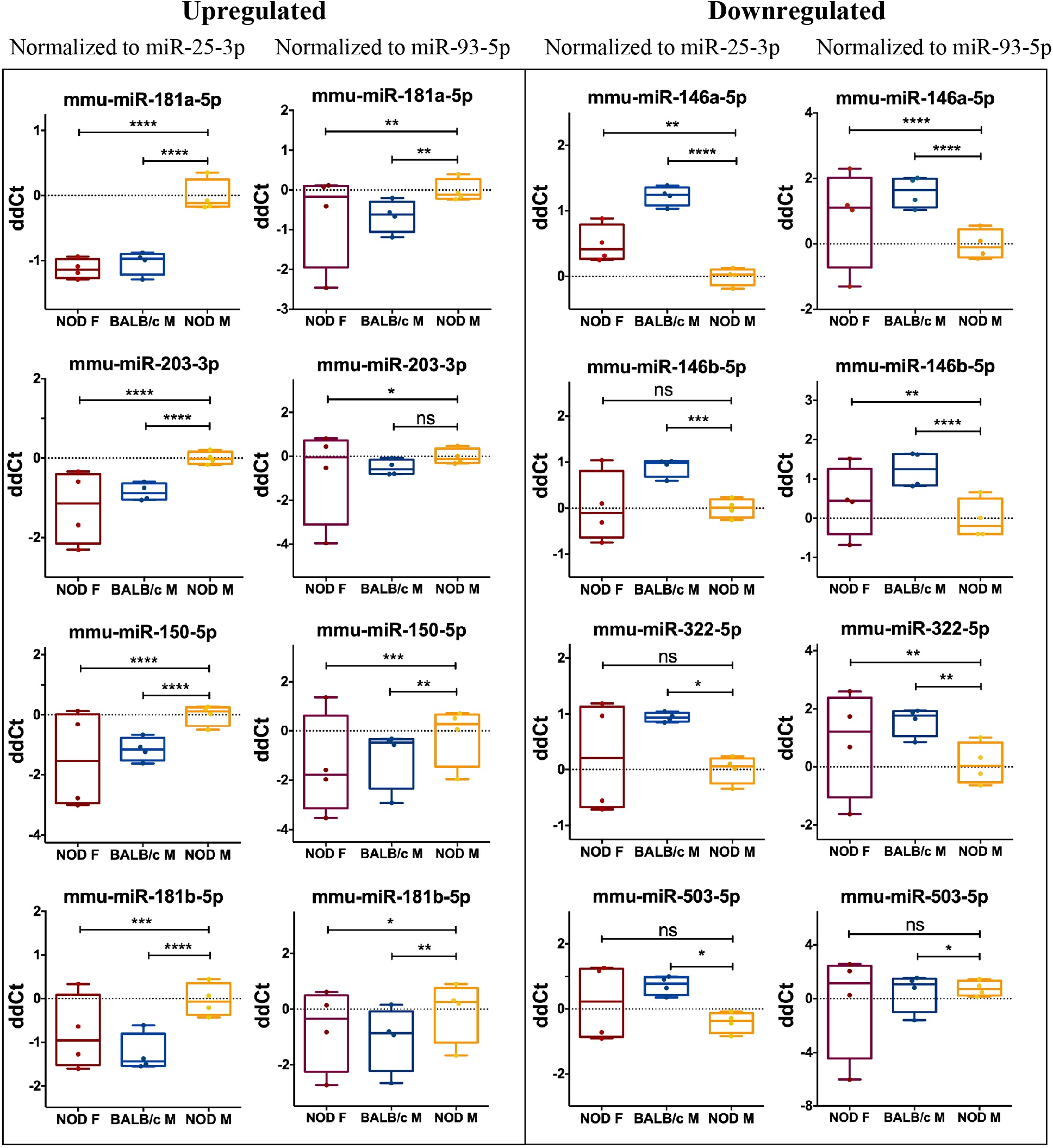
qRT-PCR validation of miRNA hits. In a separate cohort of age-matched mice, 13 miRNA hits from the sequencing data set were tested by qPCR to confirm the initial observation. Data was first normalized for plate to plate variation using a spike-in RNA and then to two internal controls, miR-93-3p and miR-25-3p. Of the 13 original hits, 8 miRNA were confirmed as differentially expressed. Statistics were performed on average ΔCt values; data are plotted as ΔΔCt. qPCR was performed with N=4 samples per strain, n=3 mice per sample per strain, with 3 technical replicates per sample. (* p < 0.05, ** p < 0.01, *** p < 10^−3^, **** p < 10^−4^, Kruskal-Wallis ANOVA in Graph-Pad Prism)

### qPCR confirms differential expression of multiple identified miRNAs in tears from a validation cohort

To validate the observed differential expression of these miRNAs in mouse tears, we collected pooled stimulated tears from a separate mouse cohort for analysis of our tear miRNA hits. As with the original discovery cohort, the male NOD mice in this validation cohort had notable autoimmune dacryoadenitis, while the female NOD and male BALB/c mice did not (**Figure 1E**). From the NGS data analysis, we observed that five miRNAs (miR-25-3p, miR-93-5p, miR-16-5p, miR-26a-3p and miR-23a-3p) were unchanged between the three strains (**Supplemental Fig S3A, S3B**). These miRNAs have been previously identified as endogenous controls in previous miRNA research(29). Additionally, qPCR validation on the same RNA samples showed that miR-93-3p and miR-25-5p were unchanged between the three strains (**Supplemental Fig S3C, S3D**). Therefore, for our validation study, in addition to the spike-in controls, UniSp3 and UniSp6, we used these two miRNAs as endogenous controls and normalized the qPCR data to them in parallel. ‘Hits’ were considered validated if they were significantly different with a fold change in the same direction in male NOD samples compared to both female NOD and male BALB/c samples and when normalized to at least one of the two endogenous miRNA controls.

These assays showed that miRNAs miR-146a-5p, miR-146b-5p, miR-322-5p and miR-503-3p were significantly downregulated, while miR-181a-5p, miR-181b-5p, miR-203-3p and miR-150-5p were significantly upregulated in male NOD mice tears as compared to tears of male BALB/c and female NOD (**Figure 4**). However, miR-146b-5p and miR-322-5p were significantly downregulated in both comparisons when normalized to only miR-93-3p, while miR-203-3p was upregulated in both comparisons when normalized to only miR-25-3p (**Table 2**).

**Figure 4.**
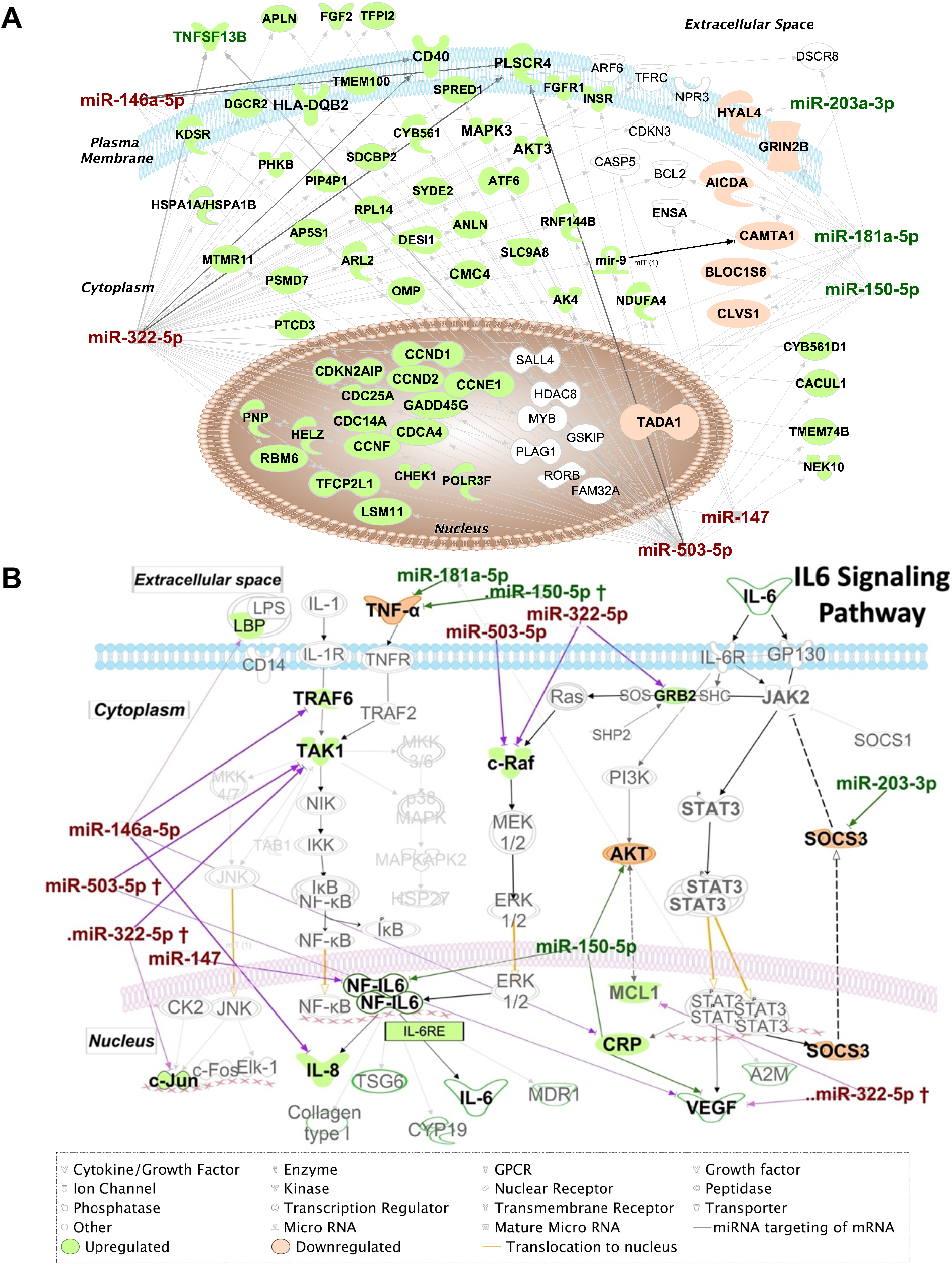
IPA Network analysis of targeted miRNAs. (A) IPA Network map of miRNA hits and their mRNA targets. Genes targeted by >1 miRNA hits are shown according to their subcellular localization. Arrows represent post-transcriptional silencing of genes by connected miRNA. Downregulated miRNA hits are shown in red, upregulated ones are in green, whereas gene targets likely to be upregulated are shaded in green, and those likely to be downregulated are shaded in orange. Gene icons shaded in white are targeted by both up and downregulated miRNA. (B) Pathway analysis results suggest upregulation of IL-6 family cytokines and those transcribed through the IL-6RE response element. In the NF-κB signaling, TRAF6 and TAK1 are targets of hits miR-146a-5p and miR-322-5p. Downregulation of these hits may lead to upregulation of TRAF6 and TAK1 and increase the transcription of the IL6 Response element (IL6RE), resulting in increased mRNA levels of cytokines. Of these, IL8 is directly targeted by miR-146a-5p and its levels could be particularly upregulated with depletion of this miRNA. Also of key import is the targeting of SOCS3 by the upregulated miRNA miR-203-3p. SOCS3 is a negative regulator of the JAK2/STAT3 pathway and is required to turn off the pathway to prevent excessive production of cytokines.

**Table 2.**
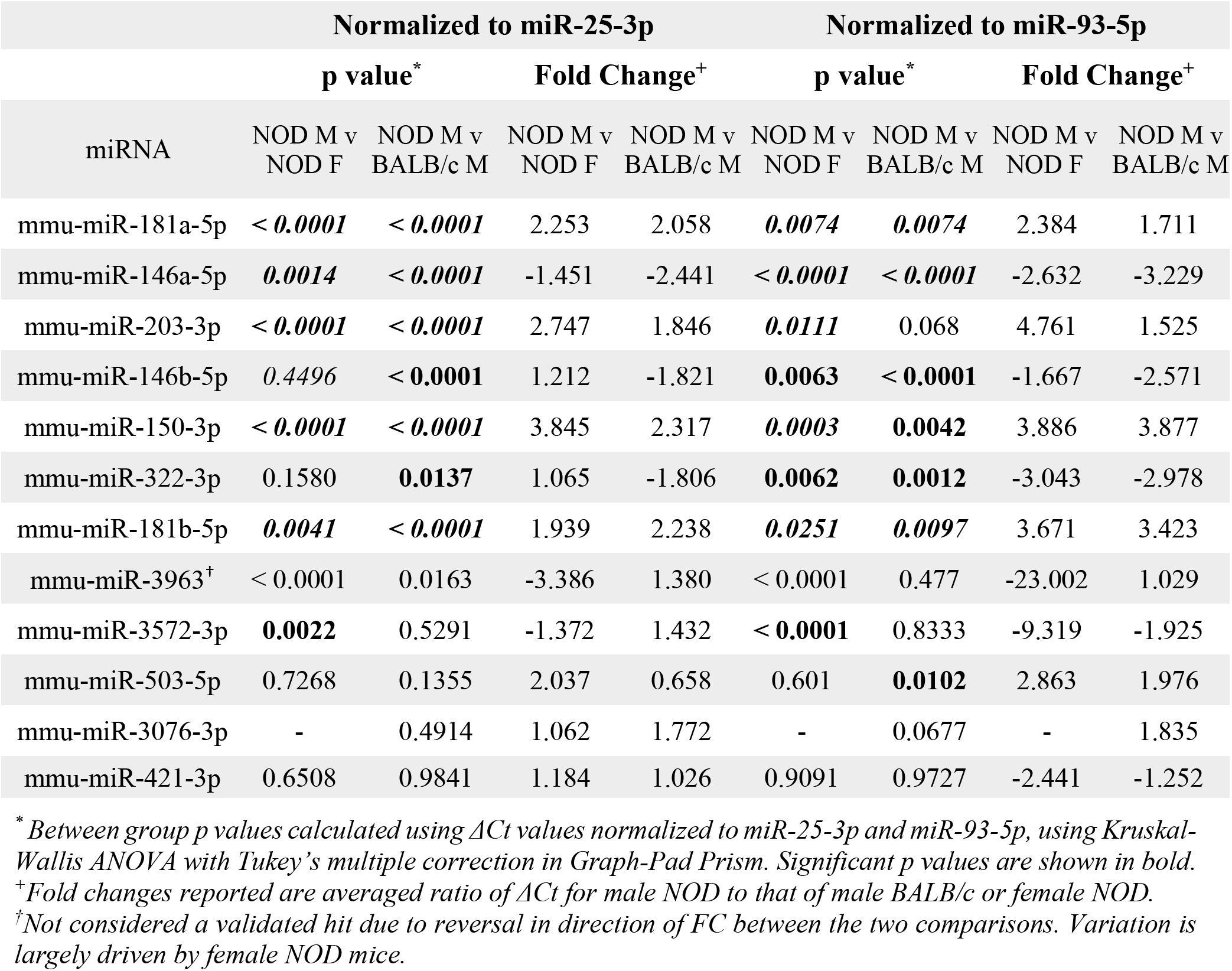
qPCR validation of differentially expressed miRNA in male NOD mice tears.

Of the hits we could not validate, the miR-147-3p primer exhibited multiple melt curves across all samples for all three replicates and did not meet the quality control requirements for qPCR. miR-3076-3p also did not meet quality control requirements due to failed amplification in 2 out of 4 samples of female NOD and male BALB/c mouse tears, and therefore lacked power for statistical analysis. Differential expression of three other miRNAs, miR-3572-5p, miR-421-3p, miR-3963 could not be validated by qPCR (**Supplemental Fig S4**).

### IPA identifies dysregulation of molecular functions and immunomodulation

To understand the pathways and cellular processes involving the qPCR-validated dysregulated miRNAs, we used Ingenuity Pathway Analysis (IPA). IPA identified mRNA targets of hits that are a) experimentally validated or b) predicted targets with high probability scores (from the TargetScan database). Biological processes with a relatively higher number of such mRNA targets are more likely to be dysregulated by altered expression of miRNA hits and are, therefore, termed ‘overrepresented’. **Supplemental Fig S5** shows the processes most likely to be affected, in decreasing order of statistical probability. The most overrepresented cellular processes and functions were differentiation and hematopoiesis of mononuclear leukocytes (p<10^−55^), leukopoiesis (p<10^−53^) and lymphopoiesis (p<10^−48^), followed by connective tissue cells (p <10^−42^), lymphatic system cells (p<10^−42^) proliferation of mononuclear leukocytes (p<10^−41^) and lymphocytes (p <10^−41^). Proliferation processes related to other cell types, such as those of liver (p<10^−16^), kidney (p<10^−12^), and heart (p<10^−6^) were relatively underrepresented. Among cellular processes leading to disease development, cancers such as nonhematologic malignant neoplasm and head and neck tumor was the most over-represented (p<10^−43^).

**Figure 5A** shows mRNAs targeted by more than one miRNA hit, shown within the context of the subcellular location of the encoded protein. The mRNA targets include those predicted to be targeted by miRNA hits with high probability based on in-silico seed region matching, and targets that are experimentally validated. Several mRNA targets appear to be transcription factors implicated in cell cycle progression such as CDKN2AIP, CDC25A, CDCA4, CDC14, CCDN1 and CCDN2. Also, of interest is PLSCR4 (Calcium dependent Phospholipid scramblase 4), an ATP-independent, transmembrane lipid transporter responsible for inducing non-specific bidirectional movement of phospholipids in the cell membrane during cell activation that has also been proposed as a potential mediator of phosphatidylserine in the outer leaflet of the plasma membrane in apoptosis(30). PLSCR4 is targeted by three of the downregulated miRNA hits, miR-146a-5p, miR-322-5p and miR-503-5p. Genes targeted by miRNA hits known to be involved in immunological processes include CD40, TNFSF13B and HLA-DBQ2, with their expression predicted to increase due to the observed downregulation of their regulating miRNAs. Interaction of CD40 with its ligand CD40L (or CD154) plays an important role in immunity as it promotes plasma cell differentiation, antibody production and is required for optimal immune response to most antigens. This is in line with previous studies demonstrating that CD40 is overexpressed in SG ductal epithelial cells, lymphocytes, and endothelial cells in SS patients(31). Antibody blockade of CD40 has also shown therapeutic potential for treatment of SS in clinical trials(32).

Several key proinflammatory mediators are also targets of dysregulated miRNA. To illustrate this, IPA results focused solely on IL6 signaling pathways are shown in **Figure 5B**. IL8 and C-reactive protein (CRP) mRNAs are targets of the downregulated miRNA hits, miR-322-3p and miR-146a-5p. TAK1 and TRAF6 of the NF-kB pathway and c-Raf of the MAPK/ERK pathway are also targets of the downregulated hits, miR-322-5p, miR-146a-5p and miR-503-5p. This downregulation may lead to upregulation of IL6-Response elements (**Figure 5B**) promoting upregulation of cytokines such as IL6 and IL8 which are highly relevant to SS pathogenesis and upregulated in salivary glands(33), serum and tears(34) of SS patients. SOCS3, a negative regulator of the JAK2/STAT3 pathway, is targeted by overexpressed miR-203-3p, which may result in its downregulation leading to the constitutive transcription of CRP, VEGF and A2M (**Figure 5B**). SOCS3 downregulation may lead to enhanced activity of pro-inflammatory IL6 family cytokines through its reduced ability to interaction with gp130(35) to modulate IL6 signaling, also potentially contributing to SS pathogenesis.

## DISCUSSION

Dry eye disease is a multifactorial condition linked to chronic inflammatory diseases, environmental challenges, hormonal status and even medication, However, SS-associated dry eye, which occurs through a distinct mechanism involving development of autoimmune inflammation and loss of secretory function of the LG, is arguably the most debilitating. In the absence of a definitive diagnosis and without clinical intervention, SS patients with dry eye symptoms can develop corneal lesions which can require corneal transplantation to restore vision(36, 37). Unchecked autoimmune inflammation in exocrine tissues including LG can lead to formation of ectopic germinal centers which may also contribute to autoantibody production and other mediators of extraglandular symptoms of SS(38, 39). Clinical diagnosis of dry eye largely evaluates functional measures such as tear flow, tear osmolarity, tear break up time and ocular surface integrity; there are no diagnostic tests to directly assess inflammation of the LG. While the LG cannot be biopsied owing to the substantial risk of damage, tear analysis can be conducted non-invasively. Soluble constituents of tears are produced and secreted largely by the LG; thus, any damage, immune infiltration, deposition, or fibrosis in the LG should be reflected by changes in the tear composition.

Addition of an analytical tear test to the ACR-EULAR criteria that could inform clinicians on the status of LG inflammation distinguish SS patients with dry eye disease from patients with dry eye due to other causes. Tears can be collected non-invasively using Schirmer’s strips(40), obtained as part of a common clinical test for dry eye. Dysregulated miRNAs can be measured at any time after tear collections are flash-frozen and can be multiplexed to detect multiple species, as they are chemically stable when stored at −80 °C(41). A tear-based test would add significantly to the assessment of the involvement of different organs in the diagnosis of SS by other specialists. Although symptoms of dry eye disease are a part of the ACR-EULAR criteria for SS diagnosis, these are weighted to emphasize SG disease more heavily since they include an assessment of both oral symptoms and a minor SG biopsy. SS patients report uneven manifestation of ocular versus oral symptoms(11), suggesting that exocrinopathies of the LG and SG may not develop in parallel. Murine models of SS also manifest different patterns of LG and SG exocrinopathy(22, 42). Thus, diagnostic tests that emphasize identification of SG involvement may not distinguish SS patients if they experience a more dominant LG disease.

Tears are a viable source for identification of unique miRNAs denoting disease. Human tears are reported to contain over 600 distinct proteins(43) and over 300 miRNAs(17). Tears also have a higher concentration of miRNA than does urine, CSF or plasma(17). Tear miRNAs are thought to play important roles in maintaining the health of the ocular surface and mediating immune response during infection. Recently, tears have been investigated for miRNA biomarkers for Alzheimer’s diseases(15) and primary open-angle glaucoma(18) with the identification of hits showing very high sensitivity and specificity (AUC = 0.93)(18). These findings collectively suggest that changes in the tear miRNAome can reflect disease status. To our knowledge, the tear miRNAome in SS has not been investigated.

Our study has identified >550 distinct miRNAs in NOD mouse tears, substantially more than reports from previous studies investigating tear miRNAs(44). The larger pool of tear miRNAs reported here may be due to our use of the NGS sequencing approach. NGS provides a broader and more comprehensive genome readout of RNAs than do microarrays. Only 2 of our 8 validated miRNA hits, mmu-miR-146a/b-5p, have previously been linked to ocular symptoms of SS while 4 of 8 of our validated miRNAs hits including miR-181b-5p, miR-203-3p, miR-322-3p and miR-503-5p have never been previously linked to SS. Nearly 70% of human miRNA have mouse orthologs with conserved seed-region sequences(13). In our study, 6 of the 8 validated miRNA hits–mmu-miR-146a-5p, mmu-miR-146b-5p, mmu-miR-150-5p, mmu-miR-181a-5p, mmu-miR-181b-5p, and mmu-miR-203-3p – are completely identical between human and mouse(45). This highlights the utility of miRNA research in murine models of disease in studying development of SS. Of the identified miRNA that are not completely identical between human and mouse, miR-503-5p differs from its human counterpart (hsa-miR-503-5p) by 1 nucleotide while identified orthologs are known for miR-322-5p (hsa-miR-424-5p), so both miRNAs could likewise be evaluated in any analyses of SS patient tears or tissue.

miR-150-5p, 146a/b-5p, and miR-181a-5p, identified here as dysregulated in tears, have been previously linked to SS in other studies of SS. One study investigating peripheral blood mononuclear cells (PBMC) from SS, SLE and healthy control subjects found miR-150-5p to be the only miRNA downregulated in SS patients as compared to SLE and healthy controls(46). However, other studies have found miR-150-5p to be elevated in SS patients’ saliva, minor SG, and serum (**Supplemental Table S1**), consistent with the increased expression in male NOD mice tears. Altered expression of miR-146a-5p and miR-146b-5p has been reported in plasma/serum and PBMCs of SS patients as well as in saliva, SG and LG from animal models of SS (**Supplemental Table S1**). A microarray of 40 miRNAs in SS patient tears showed that miR-146a-5p was downregulated in tears of primary SS patients (i.e., SS patients who lack other autoimmune diseases(1)) relative to healthy subjects but was significantly upregulated in tears of patients with secondary SS (i.e., SS patients who also have other autoimmune diseases(1)) relative to patients with primary SS(47). In SS-prone B6DC mice, miR-146a levels were upregulated in LG and submandibular SG at 8 weeks but downregulated at 20 weeks, while its levels were increased significantly in PBMC of 20 weeks old mice. Dysregulated expression of miR-146a/b-5p has also been found in animal studies and in various biofluids in patients with SLE and RA and may serve as a more general indicator of autoimmunity. A more comprehensive analysis of the changes in miR-146a/b-5p over time in tears, in parallel with a more comprehensive analysis of LG disease status, may be necessary to understand its utility in diagnosis of SS.

Dysregulation of miR-181a-5p has also been described in SS, with several studies reporting its upregulation in PBMC of SS patients relative to healthy subjects. In SG of SS patients, this miRNA was downregulated relative to healthy controls, but within the SS patient cohort it was upregulated in the SG of those exhibiting more profound decreases in salivary flow and high SG focus scores(48). Intriguingly, this miRNA is also implicated in regulation of the differentiation of germinal center B cells(49), which may also be found in the ectopic germinal centers that form at late stages of SS in exocrine glands.

All validated and dysregulated miRNAs identified here are expressed in blood cells and function in immune cell development and differentiation. Unsurprisingly, pathway analysis of the miRNA hits identifying likely gene targets and the biological processes involving these gene products include differentiation, maturation and proliferation of T and B lymphocytes. Tears produced by the LG and other ocular surface tissues drain through the canaliculus into nasolacrimal ducts. Here, tear components including miRNAs can be reabsorbed into the blood vessels surrounding the cavernous body of the nasolacrimal ducts(19) and shuttled back to the LG. These tear constituents may have access to draining lymph nodes through the tear duct associated lymphoid tissue (TALT). We hypothesize that a primary function of tear miRNAs is the regulation and homeostasis of local immune responses in the LG and ocular surface; dysregulated miRNAs may therefore contribute to induction and perpetuation of autoimmune processes by priming immune cells in the TALT that migrate to the LG in SS.

Precursor miR-146a/b and miR-181a miRNAs participate in immune cell development and differentiation of T and B cells from hematopoietic stem cells(50). miR-150-5p is detected in human serum(51), while its increased expression is linked to autoimmune disease(52). Its expression is decreased in serum of patients with B cell malignancies, and in double negative thymocytes, but its levels increase in differentiating T lymphocytes(53, 54). miR-150-5p is highly expressed in naïve T and B cells, with levels decreasing in mature B cells(53). miR-150-5p is thought to maintain B cells in the quiescent stage in lymphoid organs and to regulate their expansion by targeting the transcription factor MYB(52). It is also involved in natural killer cell maturation and development(55). These findings, along with our observation of its upregulation in male NOD tears, suggest it may participate in local T and B cell maturation with its levels possibly reflecting the degree of immune cell infiltration of the LG.

Murine miR-322 promotes Th17 cell differentiation through its targeting of the transcription factor NF-kB subunit p50(56), interesting because of the prominent role of Th17 and IL-17 as disease drivers in SS(57). miR-322 may also act as an anti-inflammatory agent through its ability to suppress cytokine secretion and signaling(58). This function is consistent with our IPA analysis which highlights the targeting of master regulators of IL6 production (TAK1 and c-Raf) by miR-322-5p (**Figure 5B**). Hsa-miR-424, the human ortholog of murine miR-322, as well as miR-503-5p are also implicated in monocyte differentiation(59, 60). Suppressor of cytokine signaling 3 (SOCS3), a negative feedback regulator of the JAK2/STAT3 pathway, is targeted by upregulated miR-203-3p which may lead to its decreased expression. As a result, IL6-mediated signaling of the JAK2/STAT3 pathway may be enhanced in cells with increased miR-203-3-p.

While most of our miRNA hits have complete sequence homology and the others have known orthologs, other species-specific differences may not be translatable. The predisposition of the male and female NOD mice to development of diabetes, although at an older age than the 12–14-week aged mice used for tear collection, is a confounding factor. Animal models may also not fully represent the complexity of SS disease progression. For instance, very little is known about development of systemic manifestations of SS in most murine disease models, while features of later B-cell mediated exocrinopathy are not pronounced in the NOD model. In addition, the pooling strategy used to obtain RNA of sufficient quantity and quality may have reduced our overall sensitivity, especially for miRNAs that are lowly expressed. However, these conditions were still sufficient to detect several significantly altered miRNAs, half of which were experimentally validated in separate samples. In conclusion, we report here the identification and validation of 8 dysregulated miRNAs in tears that have potential for use in diagnosis of ocular (LG) involvement in SS. Of these, 4 have been previously identified in other biofluids or organs in SS patients or disease models, while 4 are newly implicated in SS. This list of differentially expressed tear miRNAs is first of its kind, is of high priority for future investigation in SS patient tears and may potentially distinguish ocular involvement in SS patients.

## Supporting information

Arrive Checklist

R code for statistics

## ACKNOWLEDGEMENTS

This work was supported by RO1 EY011386 to SHA. Further support for the project came from P30EY029220, and an unrestricted departmental grant from Research to Prevent Blindness (RPB), New York, NY 10022.

## AUTHOR CONTRIBUTIONS

All authors were involved in drafting the article or revising it critically for important intellectual content, and all authors approved the final version to be published.

### Study conception and design

Hjelm, Hamm-Alvarez, Singh Kakan, Edman

### Acquisition of data

Kakan, Yao.

### Analysis and interpretation of data

Kakan, Hjelm, Hamm-Alvarez, Edman, Okamoto Authors declare no conflicts of interest.

## DATA AVAILABILITY

The authors confirm that all data underlying the findings are fully available without restriction. All relevant data are within the paper, its supporting information files. The raw sequencing data will be made available online at the sequence read archive (Accession: PRJNA769738).

## Supplemental Data

**Figure S1.**
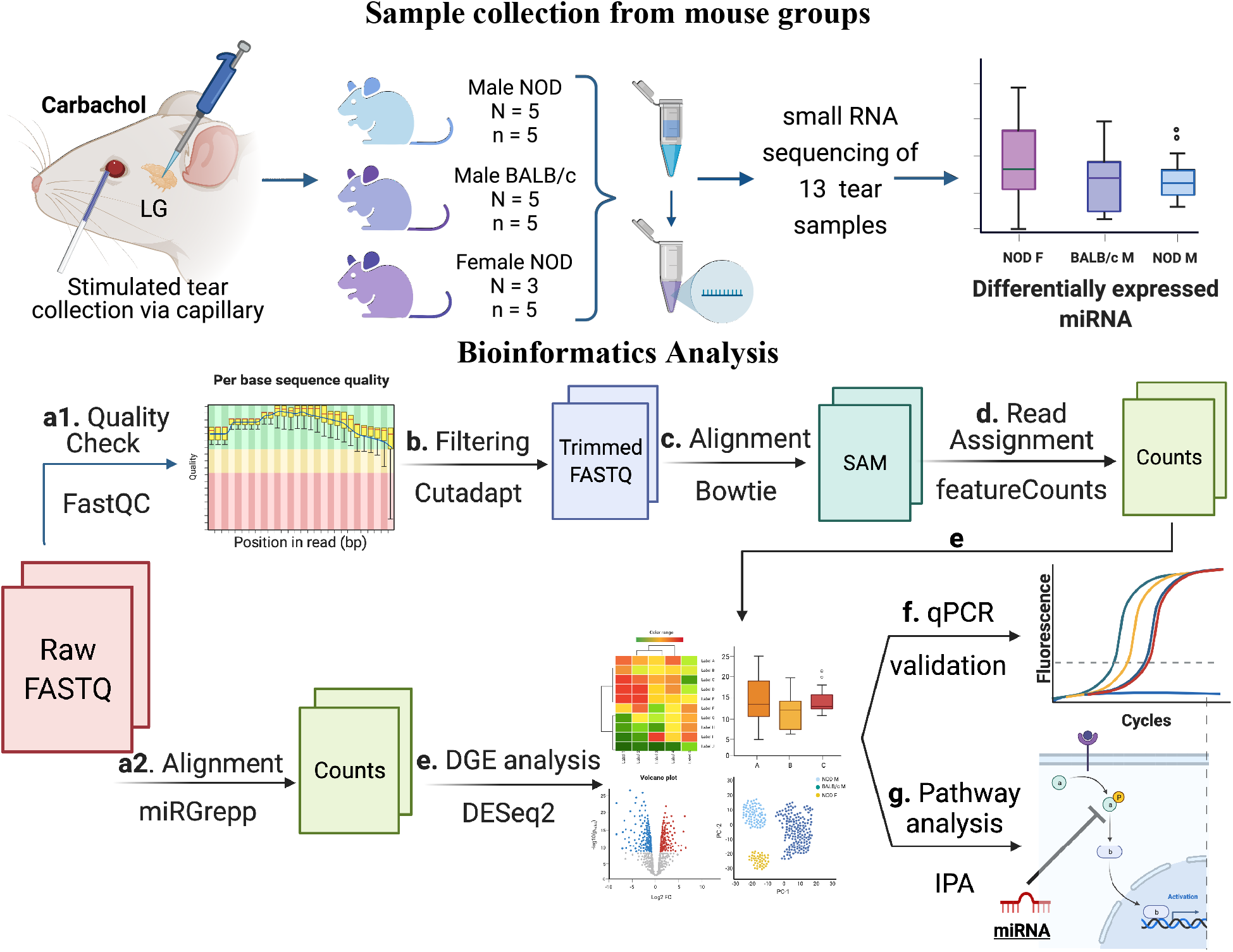
Schematic overview of experiments and data analysis. Tears were collected by stimulating both LG from each mouse with 50 μM Carbachol. Tears were pooled from 5 mice for each sample, with 5 such samples for male NOD and BALB/c mice and 3 for female NOD mice. In total, 13 samples were sent for RNA sequencing, and raw data was analyzed as follows: (a1) Quality assessment by FASTqc showed presence of 3’ adapters in the reads; (b) Using cutadap, 3’ adapters were trimmed and reads with quality scores <20 or length <15 were removed; (c) Trimmed reads were aligned to mouse genome GRCm38 using Bowtie(61); and (d) Aligned reads were annotated using featureCounts(62). Additionally, (a2) raw FASTQ files were also run through our internal miRGrepp pipeline(27) and (e) miRNA counts were normalized, and plotted to assess the quality of the data with statistics on these read counts performed using R package, DESEq2(63); (f) Shortlisted hits were validated by qPCR in a separate cohort of mice; and and (g) Pathway analysis of the hits was done in Ingenuity Pathway Analysis (Qiagen). Created with BioRender

**Figure S2.**
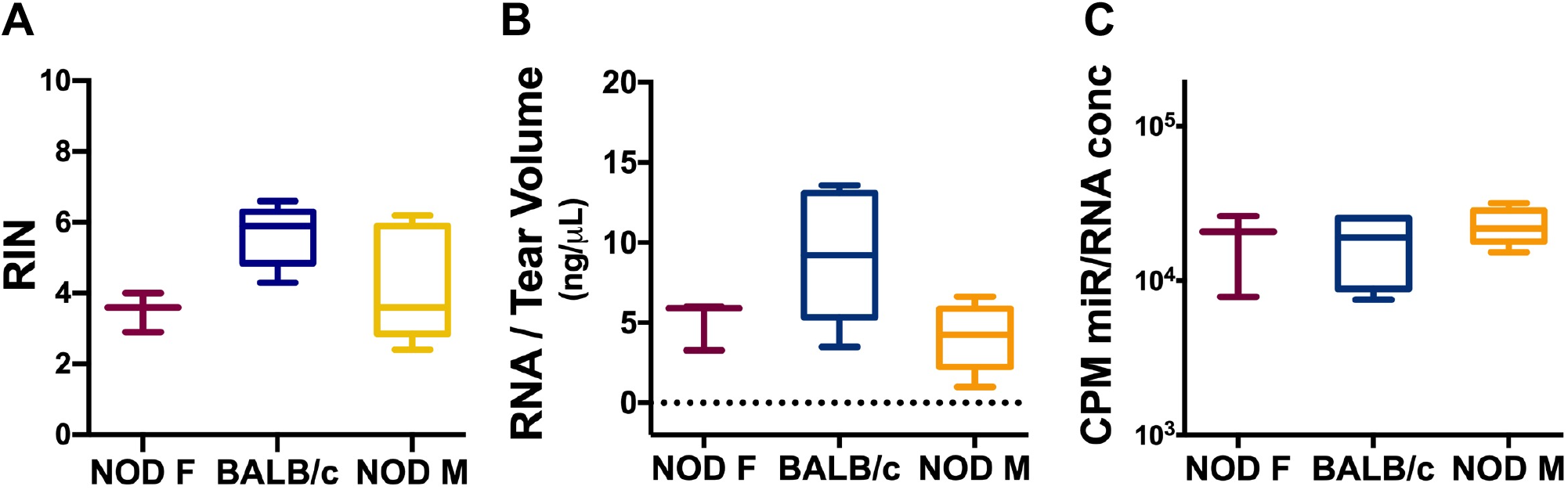
Quality assessment of RNA isolated from pooled tears of female NOD, male BALB/c and male NOD mice using Tapestation. (A) There was no significant difference in the amount of RNA isolated relative to the total tear volume isolated for each sample. (B) There was no significant difference in the RNA Integrity Number (RIN) values between samples from the three groups. (C) Counts per million (CPM) miRNA reads for each sample aligning to miRbase v22.0 did not differ significantly between the three groups. Data are plotted as boxplots showing mean with 75% to 25% IQR and whiskers show the range. N=5 samples for male NOD and BALB/c, 3 samples for female NOD; n=5 mice per sample. Samples analyzed by Kruskal-Wallis ANOVA with p<0.05 considered significantly different.

**Figure S3.**
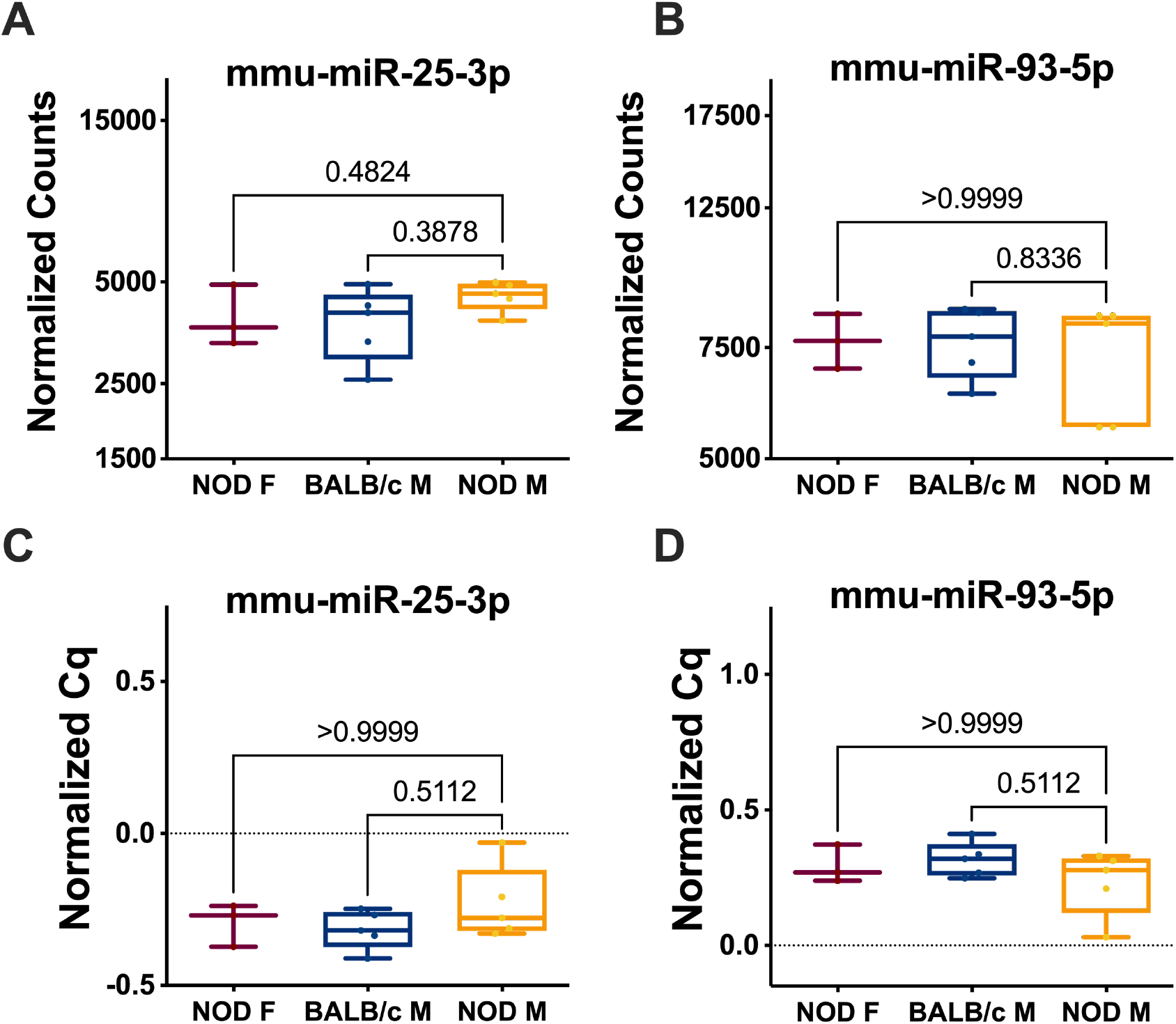
Comparison of expression of endogenous control miRNAs mmu-miR-25-3p and mmu-miR-93-5p in RNA isolated from tears of male NOD, female NOD, and male BALB/c mice. miRNA sequencing data analysis of mouse tears showed that there were no significant differences in the expression levels of (A) miR-25-3p and (B) miR-93-5p. Data for (A) and (B) are plotted as counts normalized by DESeq2 box showing all points from minimum to maximum. Each sample from the three strains had tears pooled from 5 mice, with 3 samples for the NOD female and 5 samples for male NOD and BALB/c mice. On the same set of samples, qPCR showed that miRNAs (C) miR-25-3p and (D) miR-93-5p had very similar expression levels between the three strains. Median Cq values are plotted as boxplots showing mean with 75% to 25% IQR and whiskers show the range. N=5 samples for male NOD and BALB/c, 3 samples for female NOD; n=5 mice per sample. ns - not significant, Kruskal-Wallis ANOVA, p=0.05.

**Figure S4.**
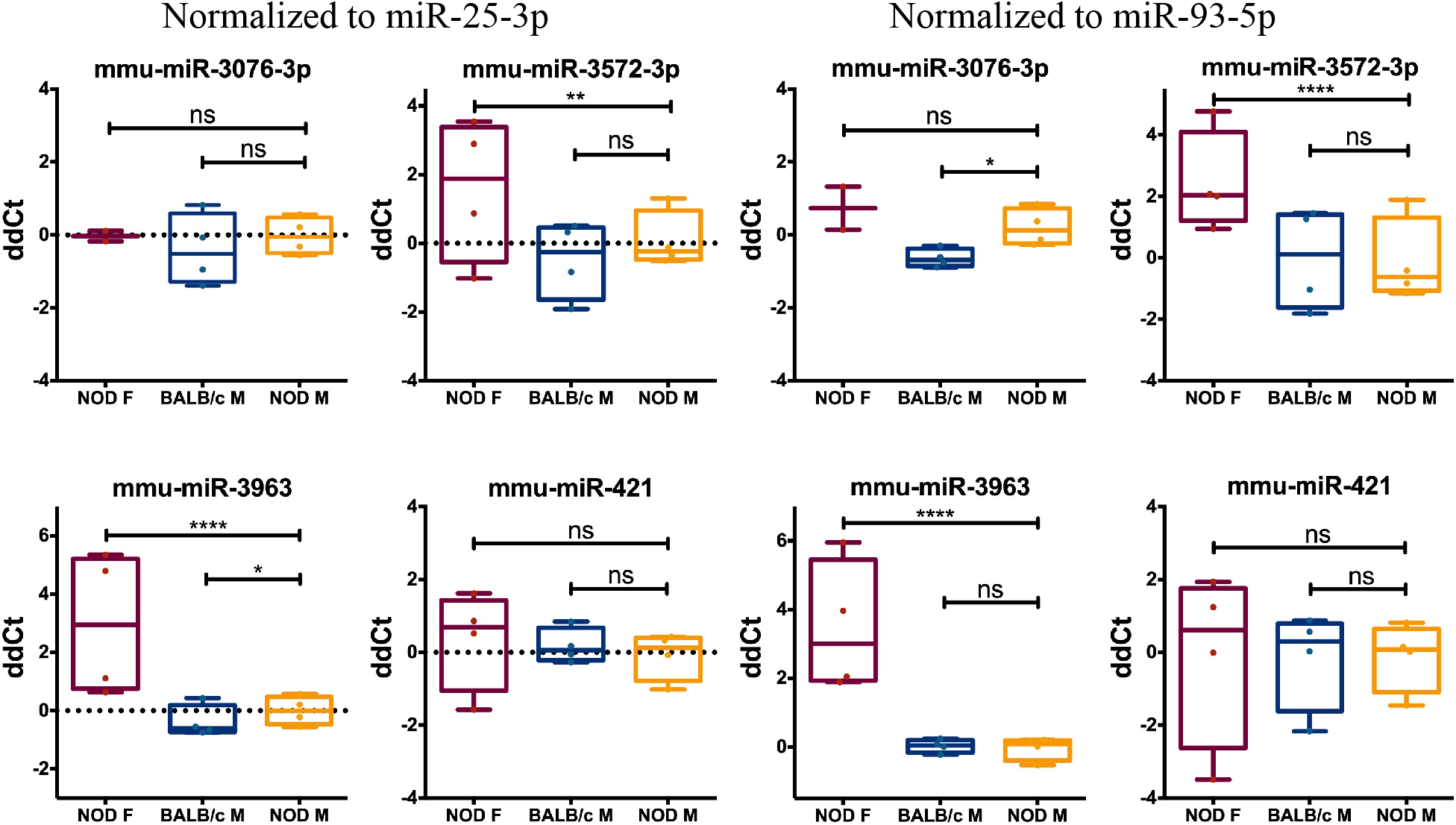
Assessment of miRNA hits predicted from the bioinformatics analysis which were not validated by qRT-PCR. Differential expression of 4 miRNAs were not confirmed in the qRT-PCR validation. miRNA miR-3076-3p expression was not significantly different from that of female NOD mice and was modestly upregulated in male NOD tears as compared to male BALB/c. miR-3572-3p was found to be downregulated in male NOD mice tears as compared to female NODs, but modestly upregulated when compared to male BALB/c. miR-421-3p was modestly downregulated in male NOD tears with respect to male BALB/c and female NOD, but this difference was not significantly different in either comparison. Mean ΔΔCt values are plotted as boxplots showing mean with 75% to 25% IQR and whiskers show the range. N=4 samples for male NOD and BALB/c, and female NOD; n=3 mice per sample. * p < 0.05, ** p < 0.01, *** p < 10^−3^, **** p < 10^−4^, ns – not; significant, Kruskal-Wallis ANOVA, p=0.05.

**Figure S5.**
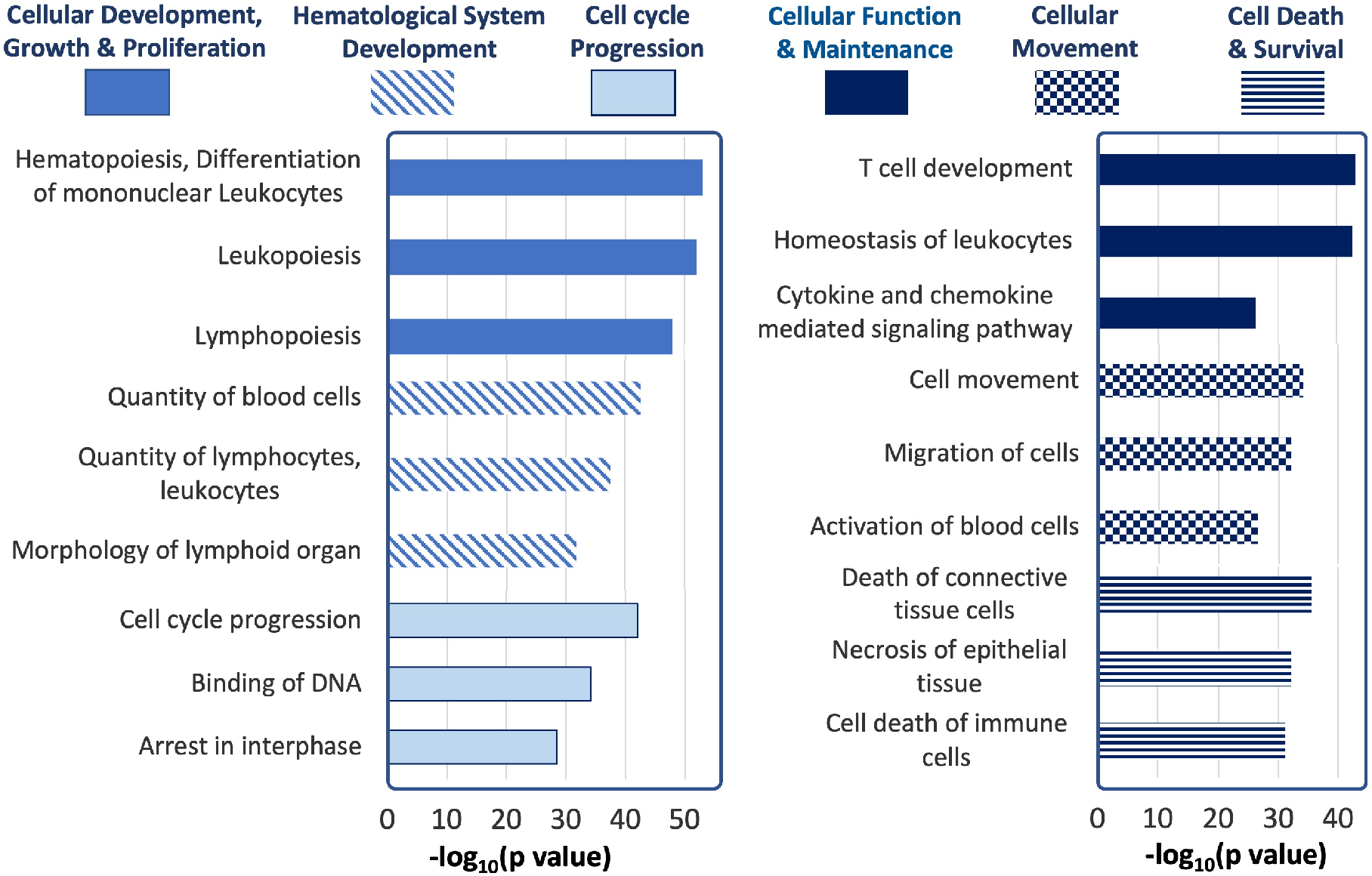
Ingenuity Pathway Analysis of tear miRNA hits. The most statistically significant biological functions and processes generated by IPA for the genes targeted by the differentially expressed tear miRNA hits, grouped by function categories (p values calculated by IPA based on **Table 1**).

**Supplemental Table S1.**
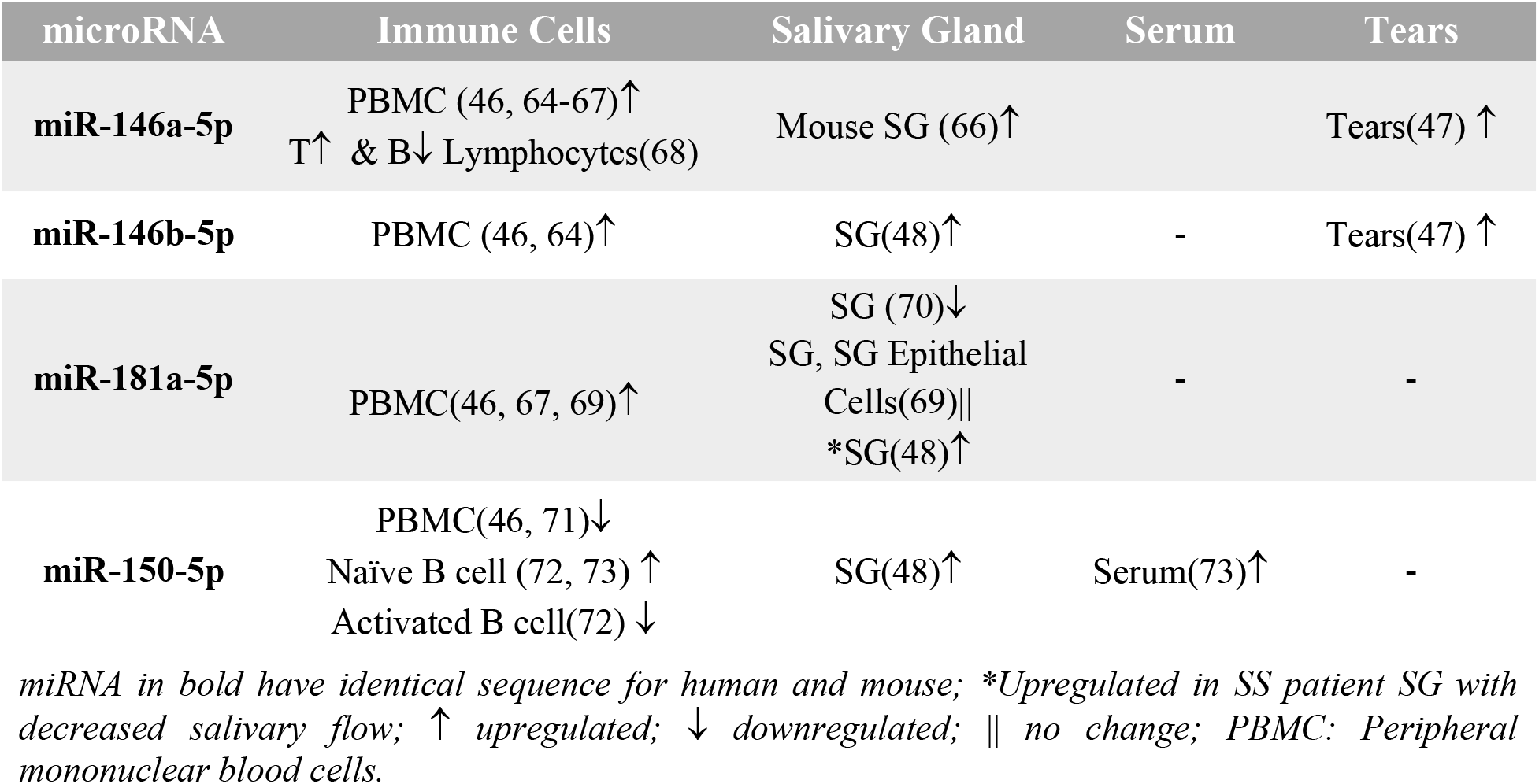
Literature summary for urevious miRNA hits in studies on SS.

## Notes

https://trace.ncbi.nlm.nih.gov/Traces/sra_sub/?subid=3569068&action=show:STUDY&acc=SRP340604&noheader=1#0

### Competing Interest Statement

The authors have declared no competing interest.

### Summary of Updates

Table 2 updated and added.

## REFERENCES

1. Vitali, C., S. Bombardieri, R. Jonsson, H. M. Moutsopoulos, E. L. Alexander, S. E. Carsons, T. E. Daniels, P. C. Fox, R. I. Fox, S. S. Kassan, S. R. Pillemer, N. Talal, and M. H. Weisman. 2002. Classification criteria for Sjögren’s syndrome: a revised version of the European criteria proposed by the American-European Consensus Group. Ann Rheum Dis 61: 554–558.

2. Price, E. J., and P. J. Venables. 2002. Dry eyes and mouth syndrome--a subgroup of patients presenting with sicca symptoms. Rheumatology (Oxford) 41: 416–422.

3. Shiboski, C. H., S. C. Shiboski, R. Seror, L. A. Criswell, M. Labetoulle, T. M. Lietman, A. Rasmussen, H. Scofield, C. Vitali, S. J. Bowman, X. Mariette, and G. International Sjogren’s Syndrome Criteria Working. 2017. 2016 American College of Rheumatology/European League Against Rheumatism Classification Criteria for Primary Sjogren’s Syndrome: A Consensus and Data-Driven Methodology Involving Three International Patient Cohorts. Arthritis Rheumatol 69: 35–45.

4. Cornec, D., V. Devauchelle-Pensec, X. Mariette, S. Jousse-Joulin, J. M. Berthelot, A. Perdriger, X. Puéchal, V. Le Guern, J. Sibilia, J. E. Gottenberg, L. Chiche, E. Hachulla, P. Yves Hatron, V. Goeb, G. Hayem, J. Morel, C. Zarnitsky, J. J. Dubost, P. Saliou, J. O. Pers, R. Seror, and A. Saraux. 2017. Severe Health-Related Quality of Life Impairment in Active Primary Sjögren’s Syndrome and Patient-Reported Outcomes: Data From a Large Therapeutic Trial. Arthritis Care Res (Hoboken) 69: 528–535.

5. Kassan, S. S., T. L. Thomas, H. M. Moutsopoulos, R. Hoover, R. P. Kimberly, D. R. Budman, J. Costa, J. L. Decker, and T. M. Chused. 1978. Increased risk of lymphoma in sicca syndrome. Ann Intern Med 89: 888–892.

6. Zufferey, P., O. C. Meyer, M. Grossin, and M. F. Kahn. 1995. Primary Sjogren’s syndrome (SS) and malignant lymphoma. A retrospective cohort study of 55 patients with SS. Scand J Rheumatol 24: 342–345.

7. Radfar, L., D. E. Kleiner, P. C. Fox, and S. R. Pillemer. 2002. Prevalence and clinical significance of lymphocytic foci in minor salivary glands of healthy volunteers. Arthritis Rheum 47: 520–524.

8. Xu, K. P., S. Katagiri, T. Takeuchi, and K. Tsubota. 1996. Biopsy of labial salivary glands and lacrimal glands in the diagnosis of Sjögren’s syndrome. J Rheumatol 23: 76–82.

9. Pavlidis, N. A., J. Karsh, and H. M. Moutsopoulos. 1982. The clinical picture of primary Sjogren’s syndrome: a retrospective study. J Rheumatol 9: 685–690.

10. Zhang, Y., T. Lin, A. Jiang, N. Zhao, and L. Gong. 2016. Vision-related quality of life and psychological status in Chinese women with Sjogren’s syndrome dry eye: a casecontrol study. BMC Womens Health 16: 75.

11. Meijer, J. M., P. M. Meiners, J. J. Huddleston Slater, F. K. Spijkervet, C. G. Kallenberg, A. Vissink, and H. Bootsma. 2009. Health-related quality of life, employment and disability in patients with Sjogren’s syndrome. Rheumatology (Oxford) 48: 1077–1082.

12. Helmick, C. G., D. T. Felson, R. C. Lawrence, S. Gabriel, R. Hirsch, C. K. Kwoh, M. H. Liang, H. M. Kremers, M. D. Mayes, P. A. Merkel, S. R. Pillemer, J. D. Reveille, and J. H. Stone. 2008. Estimates of the prevalence of arthritis and other rheumatic conditions in the United States. Part I. Arthritis Rheum 58: 15–25.

13. Warnefors, M., A. Liechti, J. Halbert, D. Valloton, and H. Kaessmann. 2014. Conserved microRNA editing in mammalian evolution, development and disease. Genome Biol 15: R83.

14. Gui, Y., H. Liu, L. Zhang, W. Lv, and X. Hu. 2015. Altered microRNA profiles in cerebrospinal fluid exosome in Parkinson disease and Alzheimer disease. Oncotarget 6: 37043–37053.

15. Kenny, A., E. M. Jiménez-Mateos, M. A. Zea-Sevilla, A. Rábano, P. Gili-Manzanaro, J. H. M. Prehn, D. C. Henshall, J. Ávila, T. Engel, and F. Hernández. 2019. Proteins and microRNAs are differentially expressed in tear fluid from patients with Alzheimer’s disease. Sci Rep 9: 15437.

16. Friedman, R. C., K. K. Farh, C. B. Burge, and D. P. Bartel. 2009. Most mammalian mRNAs are conserved targets of microRNAs. Genome Res 19: 92–105.

17. Weber, J. A., D. H. Baxter, S. Zhang, D. Y. Huang, K. H. Huang, M. J. Lee, D. J. Galas, and K. Wang. 2010. The microRNA spectrum in 12 body fluids. Clin Chem 56: 1733–1741.

18. Tamkovich, S., A. Grigor’eva, A. Eremina, A. Tupikin, M. Kabilov, V. Chernykh, V. Vlassov, and E. Ryabchikova. 2019. What information can be obtained from the tears of a patient with primary open angle glaucoma? Clinica chimica acta; international journal of clinical chemistry 495: 529–537.

19. Paulsen, F. P., U. Schaudig, and A. B. Thale. 2003. Drainage of tears: impact on the ocular surface and lacrimal system. Ocul Surf 1: 180–191.

20. Li, X., K. Wu, M. Edman, K. Schenke-Layland, M. MacVeigh-Aloni, S. R. Janga, B. Schulz, and S. F. Hamm-Alvarez. 2010. Increased expression of cathepsins and obesity-induced proinflammatory cytokines in lacrimal glands of male NOD mouse. Invest Ophthalmol Vis Sci 51: 5019–5029.

21. Doyle, M. E., L. Boggs, R. Attia, L. R. Cooper, D. R. Saban, C. Q. Nguyen, and A. B. Peck. 2007. Autoimmune dacryoadenitis of NOD/LtJ mice and its subsequent effects on tear protein composition. Am J Pathol 171: 1224–1236.

22. Hunger, R. E., C. Carnaud, I. Vogt, and C. Mueller. 1998. Male gonadal environment paradoxically promotes dacryoadenitis in nonobese diabetic mice. The Journal of clinical investigation 101: 1300–1309.

23. Zoukhri, D., R. R. Hodges, D. Byon, and C. L. Kublin. 2002. Role of proinflammatory cytokines in the impaired lacrimation associated with autoimmune xerophthalmia. Invest Ophthalmol Vis Sci 43: 1429–1436.

24. Lavoie, T. N., B. H. Lee, and C. Q. Nguyen. 2011. Current concepts: mouse models of Sjogren’s syndrome. J Biomed Biotechnol 2011: 549107.

25. Council, N. R. 2011. Guide for the Care and Use of Laboratory Animals: Eighth Edition. The National Academies Press, Washington, DC.

26. Abramoff, M. D., P. J. Magalhães, and S. J. Ram. 2004. Image processing with ImageJ. Biophotonics international 11: 36 – null.

27. Kakan, S. S. 2019. miRGrep. Github.

28. Hamm-Alvarez, S. F., S. R. Janga, M. C. Edman, S. Madrigal, M. Shah, S. E. Frousiakis, K. Renduchintala, J. Zhu, S. Bricel, K. Silka, D. Bach, M. Heur, S. Christianakis, D. G. Arkfeld, J. Irvine, W. J. Mack, and W. Stohl. 2014. Tear cathepsin S as a candidate biomarker for Sjogren’s syndrome. Arthritis Rheumatol 66: 1872–1881.

29. Schwarzenbach, H., A. M. da Silva, G. Calin, and K. Pantel. 2015. Data Normalization Strategies for MicroRNA Quantification. Clin Chem 61: 1333–1342.

30. Frasch, S. C., P. M. Henson, J. M. Kailey, D. A. Richter, M. S. Janes, V. A. Fadok, and D. L. Bratton. 2000. Regulation of phospholipid scramblase activity during apoptosis and cell activation by protein kinase Cdelta. J Biol Chem 275: 23065–23073.

31. Dimitriou, I. D., E. K. Kapsogeorgou, H. M. Moutsopoulos, and M. N. Manoussakis. 2002. CD40 on salivary gland epithelial cells: high constitutive expression by cultured cells from Sjögren’s syndrome patients indicating their intrinsic activation. Clin Exp Immunol 127: 386–392.

32. Benjamin A Fisher, Antonia Szanto, M. B. Wan-Fai Ng, Maximilian G Posch, Athena S Papas, Arwa M Farag, Thomas Daikeler, Bettina Bannert, Diego Kyburz, Alan J Kivitz, Steven E Carsons, David A Isenberg, Francesca Barone, Simon J Bowman, Pascal Espié, David Floch, Cyrielle Dupuy, Xiaohui Ren, Petra M Faerber, Andrew M Wright, Hans-Ulrich Hockey, Michael Rotte, Julie Milojevic, Alexandre Avrameas, Marie-Anne Valentin, James S Rush, and Peter Gergely. 2020. Assessment of the anti-CD40 antibody iscalimab in patients with primary Sjögren’s syndrome: a multicentre, randomised, double-blind, placebo-controlled, proof-of-concept study. The Lancet Rheumatology 2: 10.

33. Yoshimoto, K., T. Fujimoto, A. Itaya-Hironaka, T. Miyaoka, S. Sakuramoto-Tsuchida, A. Yamauchi, M. Takeda, T. Kasai, K. Nakagawara, A. Nonomura, and S. Takasawa. 2013. Involvement of autoimmunity to REG, a regeneration factor, in patients with primary Sjögren’s syndrome. Clin Exp Immunol 174: 1–9.

34. Hwang, J., S. H. Chung, S. Jeon, S. K. Kwok, S. H. Park, and M. S. Kim. 2014. Comparison of clinical efficacies of autologous serum eye drops in patients with primary and secondary Sjögren syndrome. Cornea 33: 663–667.

35. Schmitz, J., M. Weissenbach, S. Haan, P. C. Heinrich, and F. Schaper. 2000. SOCS3 exerts its inhibitory function on interleukin-6 signal transduction through the SHP2 recruitment site of gp130. J Biol Chem 275: 12848–12856.

36. Vivino, F. B., P. Minerva, C. H. Huang, and S. E. Orlin. 2001. Corneal melt as the initial presentation of primary Sjögren’s syndrome. J Rheumatol 28: 379–382.

37. Murtagh, P., R. Comer, and G. Fahy. 2018. Corneal perforation in undiagnosed Sjögren’s syndrome following topical NSAID and steroid drops post routine cataract extraction. BMJ Case Rep 2018.

38. Martin-Nares, E., and G. Hernandez-Molina. 2019. Novel autoantibodies in Sjogren’s syndrome: A comprehensive review. Autoimmun Rev 18: 192–198.

39. Mahmoud, T. I., J. Wang, J. L. Karnell, Q. Wang, S. Wang, B. Naiman, P. Gross, P. Z. Brohawn, C. Morehouse, J. Aoyama, C. Wasserfall, L. Carter, M. A. Atkinson, D. V. Serreze, H. Braley-Mullen, T. Mustelin, R. Kolbeck, R. Herbst, and R. Ettinger. 2016. Autoimmune manifestations in aged mice arise from early-life immune dysregulation. Sci Transl Med 8: 361ra137.

40. Hamm-Alvarez, S. F., C. T. Okamoto, S. R. Janga, D. Feigenbaum, M. C. Edman, D. Freire, M. Shah, R. Ghanshani, W. J. Mack, and M. F. Lew. 2019. Oligomeric α-synuclein is increased in basal tears of Parkinson’s patients. Biomarkers in medicine 13: 941–952.

41. Schirmer, O. 1903. Studien zur Physiologie und Pathologie der Tränenabsonderung und Tränenabfuhr. Albrecht von Graefes Archiv für Ophthalmologie 56: 197–291.

42. Tellefsen, S., M. K. Morthen, S. M. Richards, S. M. Lieberman, R. Rahimi Darabad, W. R. Kam, and D. A. Sullivan. 2018. Sex Effects on Gene Expression in Lacrimal Glands of Mouse Models of Sjögren Syndrome. Invest Ophthalmol Vis Sci 59: 5599–5614.

43. Soria, J., A. Acera, L. J. Merayo, J. A. Duran, N. Gonzalez, S. Rodriguez, N. Bistolas, S. Schumacher, F. F. Bier, H. Peter, W. Stocklein, and T. Suarez. 2017. Tear proteome analysis in ocular surface diseases using label-free LC-MS/MS and multiplexed-microarray biomarker validation. Sci Rep 7: 17478.

44. Pinazo-Durán, M. D., V. Zanón-Moreno, A. Lleó-Perez, J. J. García-Medina, C. Galbis-Estrada, M. J. Roig-Revert, C. Marco-Ramírez, M. López-Gálvez, R. Dolz-Marco, L. Duarte, C. Campos Borges, J. Salgado-Borges, and R. Gallego-Pinazo. 2016. Genetic systems for a new approach to risk of progression of diabetic retinopathy. Arch Soc Esp Oftalmol 91: 209–216.

45. Kozomara, A., M. Birgaoanu, and S. Griffiths-Jones. 2019. miRBase: from microRNA sequences to function. Nucleic Acids Res 47: D155–d162.

46. Chen, J. Q., G. Papp, S. Poliska, K. Szabo, T. Tarr, B. L. Balint, P. Szodoray, and M. Zeher. 2017. MicroRNA expression profiles identify disease-specific alterations in systemic lupus erythematosus and primary Sjogren’s syndrome. PLoS One 12: e0174585.

47. Kim, Y. J., Y. Yeon, W. J. Lee, Y. U. Shin, H. Cho, Y. K. Sung, D. R. Kim, H. W. Lim, and M. H. Kang. 2019. Comparison of MicroRNA Expression in Tears of Normal Subjects and Sjögren Syndrome Patients. Invest Ophthalmol Vis Sci 60: 4889–4895.

48. Alevizos, I., S. Alexander, R. J. Turner, and G. G. Illei. 2011. MicroRNA expression profiles as biomarkers of minor salivary gland inflammation and dysfunction in Sjögren’s syndrome. Arthritis Rheum 63: 535–544.

49. Zhu, D., C. Fang, W. He, C. Wu, X. Li, and J. Wu. 2019. MicroRNA-181a Inhibits Activated B-Cell-Like Diffuse Large B-Cell Lymphoma Progression by Repressing CARD11. J Oncol 2019: 9832956.

50. Taganov, K. D., M. P. Boldin, K. J. Chang, and D. Baltimore. 2006. NF-kappaB-dependent induction of microRNA miR-146, an inhibitor targeted to signaling proteins of innate immune responses. Proc Natl Acad Sci U S A 103: 12481–12486.

51. Punga, T., R. Le Panse, M. Andersson, F. Truffault, S. Berrih-Aknin, and A. R. Punga. 2014. Circulating miRNAs in myasthenia gravis: miR-150-5p as a new potential biomarker. Ann Clin Transl Neurol 1: 49–58.

52. Xiao, C., D. P. Calado, G. Galler, T. H. Thai, H. C. Patterson, J. Wang, N. Rajewsky, T. P. Bender, and K. Rajewsky. 2007. MiR-150 controls B cell differentiation by targeting the transcription factor c-Myb. Cell 131: 146–159.

53. Ghisi, M., A. Corradin, K. Basso, C. Frasson, V. Serafin, S. Mukherjee, L. Mussolin, K. Ruggero, L. Bonanno, A. Guffanti, G. De Bellis, G. Gerosa, G. Stellin, D. M. D’Agostino, G. Basso, V. Bronte, S. Indraccolo, A. Amadori, and P. Zanovello. 2011. Modulation of microRNA expression in human T-cell development: targeting of NOTCH3 by miR-150. Blood 117: 7053–7062.

54. Monticelli, S., K. M. Ansel, C. Xiao, N. D. Socci, A. M. Krichevsky, T. H. Thai, N. Rajewsky, D. S. Marks, C. Sander, K. Rajewsky, A. Rao, and K. S. Kosik. 2005. MicroRNA profiling of the murine hematopoietic system. Genome Biol 6: R71.

55. Bezman, N. A., T. Chakraborty, T. Bender, and L. L. Lanier. 2011. miR-150 regulates the development of NK and iNKT cells. J Exp Med 208: 2717–2731.

56. Pang, B., Y. Zhen, C. Hu, Z. Ma, S. Lin, and H. Yi. 2020. Myeloid-derived suppressor cells shift Th17/Treg ratio and promote systemic lupus erythematosus progression through arginase-1/miR-322-5p/TGF-β pathway. Clin Sci (Lond) 134: 2209–2222.

57. Lee, S. Y., S. J. Han, S. M. Nam, S. C. Yoon, J. M. Ahn, T. I. Kim, E. K. Kim, and K. Y. Seo. 2013. Analysis of tear cytokines and clinical correlations in Sjögren syndrome dry eye patients and non-Sjögren syndrome dry eye patients. Am J Ophthalmol 156: 247–253.e241.

58. Zhang, K., F. Song, X. Lu, W. Chen, C. Huang, L. Li, D. Liang, S. Cao, and H. Dai. 2017. MicroRNA-322 inhibits inflammatory cytokine expression and promotes cell proliferation in LPS-stimulated murine macrophages by targeting NF-κB1 (p50). Biosci Rep 37.

59. Forrest, A. R., M. Kanamori-Katayama, Y. Tomaru, T. Lassmann, N. Ninomiya, Y. Takahashi, M. J. de Hoon, A. Kubosaki, A. Kaiho, M. Suzuki, J. Yasuda, J. Kawai, Y. Hayashizaki, D. A. Hume, and H. Suzuki. 2010. Induction of microRNAs, mir-155, mir-222, mir-424 and mir-503, promotes monocytic differentiation through combinatorial regulation. Leukemia 24: 460–466.

60. Wang, J., G. Xiang, K. Mitchelson, and Y. Zhou. 2011. Microarray profiling of monocytic differentiation reveals miRNA-mRNA intrinsic correlation. J Cell Biochem 112: 2443–2453.

## SUPPLEMENTAL REFERENCES

61. Langmead, B., C. Trapnell, M. Pop, and S. L. Salzberg. 2009. Ultrafast and memory-efficient alignment of short DNA sequences to the human genome. Genome Biol 10: R25.

62. Liao, Y., G. K. Smyth, and W. Shi. 2014. featureCounts: an efficient general purpose program for assigning sequence reads to genomic features. Bioinformatics 30: 923–930.

63. Love, M. I., W. Huber, and S. Anders. 2014. Moderated estimation of fold change and dispersion for RNA-seq data with DESeq2. Genome Biol 15: 550.

64. Zilahi, E., T. Tarr, G. Papp, Z. Griger, S. Sipka, and M. Zeher. 2012. Increased microRNA-146a/b, TRAF6 gene and decreased IRAK1 gene expressions in the peripheral mononuclear cells of patients with Sjögren’s syndrome. Immunol Lett 141: 165–168.

65. Shi, H., L. Y. Zheng, P. Zhang, and C. Q. Yu. 2014. miR-146a and miR-155 expression in PBMCs from patients with Sjögren’s syndrome. J Oral Pathol Med 43: 792–797.

66. Pauley, K. M., C. M. Stewart, A. E. Gauna, L. C. Dupre, R. Kuklani, A. L. Chan, B. A. Pauley, W. H. Reeves, E. K. Chan, and S. Cha. 2011. Altered miR-146a expression in Sjögren’s syndrome and its functional role in innate immunity. Eur J Immunol 41: 2029–2039.

67. Peng, L., W. Ma, F. Yi, Y. J. Yang, W. Lin, H. Chen, X. Zhang, L. H. Zhang, F. Zhang, and Q. Du. 2014. MicroRNA profiling in Chinese patients with primary Sjögren syndrome reveals elevated miRNA-181a in peripheral blood mononuclear cells. J Rheumatol 41: 2208–2213.

68. Wang-Renault, S. F., S. Boudaoud, G. Nocturne, E. Roche, N. Sigrist, C. Daviaud, A. Bugge Tinggaard, V. Renault, J. F. Deleuze, X. Mariette, and J. Tost. 2018. Deregulation of microRNA expression in purified T and B lymphocytes from patients with primary Sjögren’s syndrome. Ann Rheum Dis 77: 133–140.

69. Gourzi, V. C., E. K. Kapsogeorgou, N. C. Kyriakidis, and A. G. Tzioufas. 2015. Study of microRNAs (miRNAs) that are predicted to target the autoantigens Ro/SSA and La/SSB in primary Sjögren’s Syndrome. Clin Exp Immunol 182: 14–22.

70. Wang, Y., G. Zhang, L. Zhang, M. Zhao, and H. Huang. 2018. Decreased microRNA-181a and −16 expression levels in the labial salivary glands of Sjögren syndrome patients. Exp Ther Med 15: 426–432.

71. Chen, J. Q., G. Papp, S. Póliska, K. Szabó, T. Tarr, B. L. Bálint, P. Szodoray, and M. Zeher. 2017. MicroRNA expression profiles identify disease-specific alterations in systemic lupus erythematosus and primary Sjögren’s syndrome. PLoS One 12: e0174585.

72. Iqbal, J., Y. Shen, Y. Liu, K. Fu, E. S. Jaffe, C. Liu, Z. Liu, C. M. Lachel, K. Deffenbacher, T. C. Greiner, J. M. Vose, S. Bhagavathi, L. M. Staudt, L. Rimsza, A. Rosenwald, G. Ott, J. Delabie, E. Campo, R. M. Braziel, J. R. Cook, R. R. Tubbs, R. D. Gascoyne, J. O. Armitage, D. D. Weisenburger, T. W. McKeithan, and W. C. Chan. 2012. Genome-wide miRNA profiling of mantle cell lymphoma reveals a distinct subgroup with poor prognosis. Blood 119: 4939–4948.

73. Lopes, A. P., M. R. Hillen, E. Chouri, S. L. M. Blokland, C. P. J. Bekker, A. A. Kruize, M. Rossato, J. A. G. van Roon, and T. Radstake. 2018. Circulating small non-coding RNAs reflect IFN status and B cell hyperactivity in patients with primary Sjögren’s syndrome. PLoS One 13: e0193157.

