## Supplementary material for "Tear miRNAs identified in a murine model of Sjögren’s Syndrome as potential diagnostic biomarkers and indicators of disease mechanism": Arrive Checklist

NOTE: Please save this file locally before filling in the table, DO NOT work on the file within your internet browser as changes will not be saved. Adobe Acrobat Reader (available free here) is recommended for completion.

### The ARRIVE guidelines 2.0: author checklist

#### The ARRIVE Essential 10

These items are the basic minimum to include in a manuscript. Without this information, readers and reviewers cannot assess the reliability of the findings.

| Item | | Recommendation | Section/line number, or reason for not reporting |
|---|---|---|---|
| **Study design** | 1 | For each experiment, provide brief details of study design including: | |
| | | a. The groups being compared, including control groups. If no control group has been used, the rationale should be stated. | Abstract/28-31<br>Materials and Methods/124-128 |
| | | b. The experimental unit (e.g. a single animal, litter, or cage of animals). | Materials and Methods/141-143; |
| **Sample size** | 2 | a. Specify the exact number of experimental units allocated to each group, and the total number in each experiment. Also indicate the total number of animals used. | Materials and Methods/141-148 |
| | | b. Explain how the sample size was decided. Provide details of any *a priori* sample size calculation, if done. | Materials and Methods/148-151; 182-185 |
| **Inclusion and exclusion criteria** | 3 | a. Describe any criteria used for including and excluding animals (or experimental units) during the experiment, and data points during the analysis. Specify if these criteria were established *a priori*. If no criteria were set, state this explicitly. | Materials and Methods: 128-129; 176-177 |
| | | b. For each experimental group, report any animals, experimental units or data points not included in the analysis and explain why. If there were no exclusions, state so. | Material and methods: ; 191-192 |
| | | c. For each analysis, report the exact value of *n* in each experimental group. | Figure Legends/744-748; 753-758; 764-765; Supplementary Figure S1 |
| **Randomisation** | 4 | a. State whether randomisation was used to allocate experimental units to control and treatment groups. If done, provide the method used to generate the randomisation sequence. | N/A<br>No treatments were performed in the study. |
| | | b. Describe the strategy used to minimise potential confounders such as the order of treatments and measurements, or animal/cage location. If confounders were not controlled, state this explicitly. | Materials and Methods/128-129 ; Discussion |
| **Blinding** | 5 | Describe who was aware of the group allocation at the different stages of the experiment (during the allocation, the conduct of the experiment, the outcome assessment, and the data analysis). | Materials and methods/178-179 |
| **Outcome measures** | 6 | a. Clearly define all outcome measures assessed (e.g. cell death, molecular markers, or behavioural changes). | Materials and Methods/ 140-141, 161-165, 174-176, 188-192, 203-206; Figure 1 Legend/738-740 |
| | | b. For hypothesis-testing studies, specify the primary outcome measure, i.e. the outcome measure that was used to determine the sample size. | Materials and Methods/188-192 |
| **Statistical methods** | 7 | a. Provide details of the statistical methods used for each analysis, including software used. | Materials and Methods/ 187-194; Supplementary Figure S1; Legends - Figure1/747-748, Figure2/757-758, Figure3 /765-766; Tables 1 &2 |
| | | b. Describe any methods used to assess whether the data met the assumptions of the statistical approach, and what was done if the assumptions were not met. | Materials and Methods/191-194; Supplementary Figure S1 |
| **Experimental animals** | 8 | a. Provide species-appropriate details of the animals used, including species, strain and substrain, sex, age or developmental stage, and, if relevant, weight. | Abstract/28-31;<br>Materials and methods/124-130; |
| | | b. Provide further relevant information on the provenance of animals, health/immune status, genetic modification status, genotype, and any previous procedures. | Introduction/98-110; |
| **Experimental procedures** | 9 | For each experimental group, including controls, describe the procedures in enough detail to allow others to replicate them, including: | |
| | | a. What was done, how it was done and what was used. | Materials and methods/Paragraphs 1-7 |
| | | b. When and how often. | Materials and methods/Paragraphs 1-7 |
| | | c. Where (including detail of any acclimatisation periods). | Materials and methods/Paragraphs 1-7 |
| | | d. Why (provide rationale for procedures). | Materials and methods/Paragraphs 1-7 |
| **Results** | 10 | For each experiment conducted, including independent replications, report: | |
| | | a. Summary/descriptive statistics for each experimental group, with a measure of variability where applicable (e.g. mean and SD, or median and range). | Tables 1 & 2 |
| | | b. If applicable, the effect size with a confidence interval. | N/A |

### The Recommended Set

These items complement the Essential 10 and add important context to the study. Reporting the items in both sets represents best practice.

| Item | | Recommendation | Section/line number, or reason for not reporting |
|---|---|---|---|
| **Abstract** | 11 | Provide an accurate summary of the research objectives, animal species, strain and sex, key methods, principal findings, and study conclusions. | Abstract/25-49; |
| **Background** | 12 | a. Include sufficient scientific background to understand the rationale and context for the study, and explain the experimental approach. | Introduction/53-70, 74-80, 116-120, 89-97, 111-116 |
| | | b. Explain how the animal species and model used address the scientific objectives and, where appropriate, the relevance to human biology. | Introduction/98-110, 74-84 Discussion/ |
| **Objectives** | 13 | Clearly describe the research question, research objectives and, where appropriate, specific hypotheses being tested. | Introduction/95-97; Discussion/385-393 |
| **Ethical statement** | 14 | Provide the name of the ethical review committee or equivalent that has approved the use of animals in this study, and any relevant licence or protocol numbers (if applicable). If ethical approval was not sought or granted, provide a justification. | Materials and Methods/ 130-133 |
| **Housing and husbandry** | 15 | Provide details of housing and husbandry conditions, including any environmental enrichment. | Materials and Methods/130-133 |
| **Animal care and monitoring** | 16 | a. Describe any interventions or steps taken in the experimental protocols to reduce pain, suffering and distress. | Materials and Methods: 137-139 |
| | | b. Report any expected or unexpected adverse events. | N/A |
| | | c. Describe the humane endpoints established for the study, the signs that were monitored and the frequency of monitoring. If the study did not have humane endpoints, state this. | Material and Methods/142-143 |
| **Interpretation/ scientific implications** | 17 | a. Interpret the results, taking into account the study objectives and hypotheses, current theory and other relevant studies in the literature. | Discussion/394-467 |
| | | b. Comment on the study limitations including potential sources of bias, limitations of the animal model, and imprecision associated with the results. | Discussion/468-478 |
| **Generalisability/ translation** | 18 | Comment on whether, and how, the findings of this study are likely to generalise to other species or experimental conditions, including any relevance to human biology (where appropriate). | Introduction/74-84; Discussion/400-408 |
| **Protocol registration** | 19 | Provide a statement indicating whether a protocol (including the research question, key design features, and analysis plan) was prepared before the study, and if and where this protocol was registered. | N/A |
| **Data access** | 20 | Provide a statement describing if and where study data are available. | Data availability/514-516 |
| **Declaration of interests** | 21 | a. Declare any potential conflicts of interest, including financial and non-financial. If none exist, this should be stated. | Authors declare no conflicts of interest. |
| | | b. List all funding sources (including grant identifier) and the role of the funder(s) in the design, analysis and reporting of the study. | Acknowledgments/481 -483 |
