## Supplementary material for "Tear miRNAs identified in a murine model of Sjögren’s Syndrome as potential diagnostic biomarkers and indicators of disease mechanism": R code for statistics: tear_miRNA.R

```
setwd('~/Documents/3_Parkinsons_disease/2019/Tears/2019_Grep_mir/miRGrepOut/')

library(calibrate)
library(dplyr)
library(DEGreport)
library(DESeq2)
library(DEFormats)
library(edgeR)
library(ggpubr)
library(ggsci)
library(ggplot2)
library(gridExtra)
library(pheatmap)
library(reshape2)
library(RColorBrewer)
library(scales)
library(tidyr)
library(magrittr)
library(PCAtools)
library(psych)

####### 0 Dataset ######

BALB_1 <- read.table("B1TmiR.txt", col.names = c("miR", "B1"), fill=T, as.is = 2)
BALB_1$B1 <- as.integer(BALB_1$B1)
BALB_2 <- read.table("B2TmiR.txt", col.names = c("miR", "B2"), fill=T)
BALB_3 <- read.table("B3TmiR.txt", col.names = c("miR", "B3"), fill=T)
BALB_4 <- read.table("B4TmiR.txt", col.names = c("miR", "B4"), fill=T)
BALB_5 <- read.table("B5TmiR.txt", col.names = c("miR", "B5"), fill=T)
NOD_1 <- read.table("N1TmiR.txt", col.names = c("miR", "N1"), fill=T)
NOD_2 <- read.table("N2TmiR.txt", col.names = c("miR", "N2"), fill=T)
NOD_3 <- read.table("N3TmiR.txt", col.names = c("miR", "N3"), fill=T)
NOD_4 <- read.table("N4TmiR.txt", col.names = c("miR", "N4"), fill=T)
NOD_5 <- read.table("N5TmiR.txt", col.names = c("miR", "N5"), fill=T)
NOD_F1 <- read.table("F1TmiR.txt", col.names = c("miR", "F1"), fill=T)
NOD_F2 <- read.table("F2TmiR.txt", col.names = c("miR", "F2"), fill=T)
NOD_F3 <- read.table("F3TmiR.txt", col.names = c("miR", "F3"), fill=T)

raw_count <- full_join(BALB_1, BALB_2) %>%
   full_join(BALB_3) %>%
   full_join(BALB_4) %>%
   full_join(BALB_5) %>%
   full_join(NOD_1) %>%
   full_join(NOD_2) %>%
   full_join(NOD_3) %>%
   full_join(NOD_4) %>%
   full_join(NOD_5) %>%
   full_join(NOD_F1) %>%
   full_join(NOD_F2) %>%
   full_join(NOD_F3)

rm(NOD_1, NOD_2, NOD_3, NOD_4, NOD_5, BALB_1, BALB_2, BALB_3, BALB_4, BALB_5, NOD_F1, NOD_F2, NOD_F3)

raw_count <- raw_count[-c(1:7, 1986 : 2071),]

rownames(raw_count) <- raw_count[,1]
raw_count[is.na(raw_count)] <- 0
raw_countmiR <- raw_count[,-1]
count_table = as.matrix(raw_countmiR)

#dge = DGEList(counts=count_table, group = group_)

setwd('~/Documents/3_Parkinsons_disease/2019/Tears/Grep_miRNA/')

reads = c(18106931, 15612560, 13253017,20479785,15629542, 36047838,17063228,15639822,16033798,15161811, 15323444, 17262745,15079707)
total <- colSums(count_table)
norm_counts = as.data.frame((total/reads * 100))
norm_counts$strain <- c(rep(c("B","N"), each=5), rep(c("F"), each=3))
norm_counts$strain <- factor(norm_counts$strain, levels=c("B", "N", "F"))
colnames(norm_counts) <- c("CPM", "strain")
norm_counts$RIN <- c(6.00, 5.90, 4.30, 5.40, 6.60,5.60, 3.30, 2.40,	3.60, 6.20, 2.90,3.60, 4.00)
norm_counts$amt <- c(204, 187, 22.168, 147.39, 242.24, 5.298, 143.6, 136.39, 74.63, 100.1, 44.7, 124.62, 106.34)
plot(norm_counts$CPM, norm_counts$RIN)
plot(norm_counts$CPM, norm_counts$amt)
p1 <- corr.test(norm_counts$CPM, norm_counts$amt, method="spearman")
corr.tes
ggplot(norm_counts, aes(strain, CPM, color=strain)) +
   geom_boxplot(outlier.shape = NA, width = 0.35, coef=1.8, show.legend = F, size=0.8)+
   geom_jitter(aes(fill=strain), alpha =0.7, size=1.5, show.legend = F, width=0.2) + theme_bw() +
   scale_colour_jco() +
   stat_summary(fun=mean, geom="point", size=3, color="red3", alpha = 1)

 ######## 1 One factor 3 level comparison (Set NOD as reference) ######

group_ <- c(rep(c("BALB/c", "NOD M"), each=5), rep(c("NOD F"), each=3))
sex_ <- c(rep("M", 10), rep("F", 3))
colData <- as.data.frame(cbind(c('B1', 'B2', 'B3', 'B4', 'B5', 'N1', 'N2', 'N3', 'N4', 'N5', 'F1', 'F2', 'F3'), group_, sex_))
#colData <- as.data.frame(cbind(group_, sex_))
colnames(colData) <- c('Sample', "Strain", "Sex")
rownames(colData) <- colData$Sample
colnames(count_table) <- colData$Sample

colData$Strain <- factor(colData$Strain)
colData$Strain <- relevel(colData$Strain, ref = "NOD M")
colData$Sex <- factor(colData$Sex)
colData$Sex <- relevel(colData$Sex, ref='F')

nm <- model.matrix(~Strain + Sex + Strain:Sex, data=colData)
nm_red <- model.matrix(~Strain + Sex, data=colData)

dds <- DESeqDataSetFromMatrix(
   countData = count_table,
   colData = colData,
   design = ~Strain)

dds <- dds[rowSums(counts(dds)) > 5,]

dds <- estimateSizeFactors(dds)
sizeFactors(dds)
par(mfrow=c(2,1))
boxplot(counts(dds, normalized=F), notch = F, main = " untransformed read counts ", ylab = "norm read counts")
# box plots of log2 - transformed read counts
boxplot (log2(counts(dds, normalized=T)+0.5), notch = F, main = " log2 - transformed read counts ", ylab= "log2(norm read counts)")
dim(dds)
plotMDS(counts(dds, normalized=T), col=as.numeric(group_))

#Creating the DESeq2 data object & running DESeq2

dseqNvBvS = DESeq(dds, test='LRT', reduced=nm_red)
dseqNvBvS = DESeq(dds)
#write.csv(counts(dseqNvBvS, normalized=T), file="NvBvF_tearcountsNorm.csv", row.names = T)


resNvsB = results(dseqNvBvS, name="Strain_BALB.c_vs_NOD.M", alpha=0.05)

summary(resNvsB)
sum(resNvsB$padj<0.05, na.rm=TRUE)
reSig <- resNvsB[(which(resNvsB$padj < 0.05)), ]
reSig <- reSig[order(reSig$padj),]
head(reSig)
resNvsB <- results(dseqNvBvS, tidy=TRUE, name="Strain_BALB.c_vs_NOD.M", alpha=0.05) %>%
   arrange(padj, pvalue) %>%
   tbl_df()
resNvsB
#write.csv(resNvsB, file="NvsB.csv", row.names = T)

resNvsF = results(dseqNvBvS, name="Strain_NOD.F_vs_NOD.M", alpha=0.05)
reSig <- resNvsF[(which(resNvsF$padj < 0.05)), ]
reSig <- reSig[order(reSig$padj),]
head(reSig, 20)

resNvsF <- results(dseqNvBvS, tidy=TRUE, name="Strain_NOD.F_vs_NOD.M", alpha=0.05) %>%
   arrange(padj, pvalue) %>%
   tbl_df()
resNvsF
#write.csv(resNvsF, file="NvsF.csv", row.names = F)

#subset hits

resNvB_ <- resNvsB[which(resNvsB$pvalue<0.25),] ###.x
resNvF_ <- resNvsF[which(resNvsF$pvalue<0.25),] ###.y

resNvBvF <- inner_join(resNvB_,resNvF_, by="row")
resNvBvF <- resNvBvF[,-c(4,5,8,10,11)]
resNvBvF1 <- resNvBvF[which(resNvBvF$log2FoldChange.x> 0 & resNvBvF$log2FoldChange.y>0 & resNvBvF$baseMean.x > 9),]
resNvBvF2 <- resNvBvF[which(resNvBvF$log2FoldChange.x < (-0) & resNvBvF$log2FoldChange.y< (0) & resNvBvF$baseMean.x > 9),]
resNvBvF1$log2FoldChange.x <- resNvBvF1$log2FoldChange.x * (-1)
resNvBvF1$log2FoldChange.y <- resNvBvF1$log2FoldChange.y * (-1)
resNvBvF2$log2FoldChange.x <- resNvBvF2$log2FoldChange.x * (-1)
resNvBvF2$log2FoldChange.y <- resNvBvF2$log2FoldChange.y * (-1)

#write.csv(resNvBvF1,file="down_hits.csv", row.names=T)
#write.csv(resNvBvF2,file="up_hits.csv", row.names=T)

hits <- rbind(raw_count[resNvBvF1$row,], raw_count[resNvBvF2$row,])
reads = c(18106931, 15612560, 13253017,20479785,15629542, 36047838,17063228,15639822,16033798,15161811, 15323444, 17262745,15079707)
total <- colSums(count_table)
hits_ <- (count_table/((total/reads) * 10))
#hits = as.data.frame(t(hits/total * 1000000))
#hits <- as.data.frame(t(hits/total * 1000000))
hits <- hits[-c(8:11,20:24),-1]
#hits$Strain <- factor(c(rep(c("BALB/c", "NOD"), each=5), rep(c("NODF"), each=3)), levels = c("BALB/c", "NOD", "NODF"))

hits_[rownames(hits),]

######2 DENSITY PLOTS ########

counts <- counts(dseqNvBvS, normalized=F)
design_ <- as.data.frame(colData(dseqNvBvS))
#hist((counts[,1]), main = paste("Sample" , colnames(counts))[1])

countData = as.data.frame(counts)
df_dseq = melt(countData, variable.name = "Samples", value.name = "count") # reshape the matrix 
df_dseq = data.frame(df_dseq, Condition = substr(df_dseq$Samples,1,1))
mycolors <- colorRampPalette(brewer.pal(8, "Set1"))(13)
par(mfrow=c(2,1))
plot.new()
#tiff("miR_density2.tiff", units="in", width=6, height=3.5, res=300)
ggplot(df_dseq, aes(x = log2(count+0.5), colour = Condition, fill = Samples)) + 
   geom_density(alpha = 0, size = 0.8) + 
   #facet_wrap(~Condition, ncol=3) +
   theme(legend.position = "top", axis.text=element_text(size=12), axis.title=element_text(size=16)) + 
   xlab(('log10(N counts+0.5)')) +
   theme_minimal() + xlim(-1.5,20) +
   scale_colour_manual(values=mycolors, name="") + guides(fill="none") 
dev.off()

##########3 HeatMap######

#Use unsorted res table for this
resNvB <- results(dseqNvBvS, tidy=TRUE, name="Strain_BALB.c_vs_NOD.M", alpha=0.05) %>%
   tbl_df()
resNvB

resNvF <- results(dseqNvBvS, tidy=TRUE, name="Strain_NOD.F_vs_NOD.M", alpha=0.05) %>%
   tbl_df()
resNvF

res <- full_join(resNvF, resNvB,  by='row')
res$log2FoldChange.x <- res$log2FoldChange.x * (-1)
res$log2FoldChange.y <- res$log2FoldChange.y * (-1)

rld <- rlogTransformation(dseqNvBvS)
ntd <- normTransform(dseqNvBvS)
sampleDists <- dist(t(assay(ntd)))
as.matrix( sampleDists )[1:13, 1:13]
mat = assay(ntd)[c(which(res$log2FoldChange.x >0 & res$log2FoldChange.y >0 & res$baseMean.x > 9.5 & res$pvalue.y < 0.1 & res$pvalue.x < 0.27), which(res$log2FoldChange.x < 0 & res$log2FoldChange.y < 0 & res$baseMean.x > 25 & res$pvalue.y < 0.1 & res$pvalue.x < 0.1)),]
mat = mat - rowMeans(mat)
df = as.data.frame(colData(ntd)[,c("Strain")])
colnames(df) = "Strain"
rownames(df) = colnames(mat)
my_colour = list(
   Strain=c( 'BALB/c'=pal_jco("default")(1),'NOD M'=pal_jco("default")(2)[2], 'NOD F'=pal_jco("default")(3)[3]))
tiff("tearNvBvFntd.jpeg", units="in", width=6.6, height=7.1, res=300)
pheatmap(mat, 
         annotation_col=df, 
         color = colorRampPalette(rev(brewer.pal(n = 7, name ="RdBu")))(100),
         cutree_cols = 3,
         cutree_rows = 2,
         annotation_colors = my_colour,
         annotation_legend =F,
         angle_col=45,
         angle_row =270,
         fontsize=14,
         scale='row')
dev.off()

######### 4 PCA ######

chart_design_pca <- theme(
   axis.text.x = element_text(size=14),
   axis.text.y = element_text(size=14),
   legend.text = element_text(size=12,margin = margin(t=1)),
   legend.title = element_blank(),
   axis.title.x = element_text(size=16, margin = margin(t=9, b=-4)),
   axis.title.y = element_text(size=17, margin = margin(r=5)),
   legend.position = "bottom")


dseqNvBvS <- estimateSizeFactors(dseqNvBvS)
b <- SummarizedExperiment(log2(counts(dseqNvBvS, normalized=T)+1), colData=colData)

tiff("PCA1_log2.jpeg", units="in", width=3.2, height=3.8, res=300)
plotPCA(rlog(dseqNvBvS), intgroup='Strain') + 
   labs(title = " ") +
   theme_minimal() + 
   chart_design_pca +
   ylim(-20,10) + xlim(-20,15) +
   geom_point(size=6, alpha=0.2) +
   scale_color_jco(name="Strain")
dev.off()

vsd <- assay(varianceStabilizingTransformation(dseqNvBvS))
p <- pca(vsd, metadata = colData, removeVar = 0.1)
screeplot(p, axisLabSize = 18, titleLabSize = 22)
tiff("PCAvsd.jpeg", units="in", width=4.2, height=3.4, res=300)
biplot(p, labSize = 5, pointSize = 5, gridlines.major = F, colby = "Strain",legendPosition="top")
dev.off()
pairsplot(p, pointSize = 5, gridlines.major = F, colby = "Strain",legendPosition="top")
plotloadings(p)

dds_ <- estimateSizeFactors(dds)
se <- SummarizedExperiment(log2(counts(dds_, normalized=T) + 1),colData=colData(dds_))
se <- SummarizedExperiment(log2(hits +1),colData=colData(dds_))

pc <- log2(counts(dds_, normalized=T) + 0.5) #pca of all miRNA
pc <- log2(counts(dds_, normalized=T)[rownames(hits),] + 0.5) #pca of top hits
#pc <- log2(hits_ + 1)
se <- SummarizedExperiment(pc, colData = colData(dds))
tiff("PCA_ntop.jpeg", units="in", width=3.5, height=3, res=300)
plotPCA(DESeqTransform(se), intgroup='Strain') +
   labs(title = " ") +
   theme_minimal() + 
   chart_design_pca +
   ylim(-4,4) + xlim(-5.6,5.6) +
   geom_point(size=5, alpha=0.4) +
   scale_color_jco(name="Strain")
dev.off()


####### 5 Volcano plot ######
res <- full_join(resNvsB, resNvsF, by='row')
rownames(res) <- res$row
rownames(res) <- res$row
res$log2FoldChange.x <- res$log2FoldChange.x * (-1)
res$log2FoldChange.y <- res$log2FoldChange.y * (-1)

labels <- subset(res[rownames(hits),], (pvalue.x<0.05 & abs(log2FoldChange.x)>0.5))$row
labels <- c("miR-322-3p", "miR-421-3p", "miR-146a-5p", "miR-147-3p", "miR-503-5p", "miR-146b-5p", "miR-322-5p", "3572-3p", "miR-181b-5p", "miR-203-3p", "miR-150-5p")

par(mfrow=c(2,1))
par(cex=1.3)
tiff("VolcanoNvB_2.jpeg", units="in", width=4.7, height=4.2, res=300)
with(res, plot(log2FoldChange.x, -log10(pvalue.x), xlab=expression(paste('log'[2],'FC')), ylab=expression(paste('-log'[10],' p value')), pch=19, xlim=c(-6,7.3), ylim=c(0.35,11.7), cex=0.4, cex.lab=1.35, mgp=c(2.3,1.1,0), col=scales::alpha("gray50",0.3)))
with(subset(res, abs(log2FoldChange.x)>2.5), points(log2FoldChange.x, -log10(pvalue.x), pch=19, col="orange", cex=0.75))
with(subset(res, padj.x< 0.05 & abs(log2FoldChange.x)>2.5), points(log2FoldChange.x, -log10(pvalue.x), pch=19, col="red", cex = 0.75))
with(subset(res[rownames(hits),], (pvalue.x<0.05 & abs(log2FoldChange.x)>0.5)), points(log2FoldChange.x, -log10(pvalue.x), pch=1, col=scales::alpha("blue",0.7), cex=0.9))
# Label points with the textxy function from the calibrate plot
with(subset(res[rownames(hits),], (pvalue.x<0.05 & abs(log2FoldChange.x)>0.51)), textxy(log2FoldChange.x-0.25, -log10(pvalue.x+(0.4*pvalue.x)), labs=labels, offset=0.5, m=c(-0.95,1.25), cex=0.4))
abline(h = 1.3, v = 2.5, col = "gray60", lty=3)
abline(h = 3.3, v = -2.5, col = "gray60", lty=3)
legend(-6, 11.8, legend=c(expression(paste("p"["adj"]," < 0.05 & ","log"[2],"FC < |2.5|")), expression(paste('log'[2],'FC', ' > |2.5|')),expression(paste("common hits")), "NS"), col=c("red", "orange","blue", "gray50"), pch=c(19,19,1,19), pt.cex=0.6, cex=0.6)
dev.off()


par(cex=1.3)
tiff("VolcanoNvF.jpeg", units="in", width=4.7, height=4.2, res=300)
with(res, plot(log2FoldChange.y, -log10(pvalue.y), xlab=expression(paste('log'[2],'FC')), ylab=expression(paste('-log'[10],' p value')), pch=19, xlim=c(-5,5), ylim=c(0.35,13.2) ,cex=0.4, cex.lab=1.3, mgp=c(2.2,1,0), col=scales::alpha("gray50", 0.4)))
with(subset(res, abs(log2FoldChange.y)>2.5), points(log2FoldChange.y, -log10(pvalue.y), pch=19, col="orange", cex=0.7))
with(subset(res, padj.y< 0.05 & abs(log2FoldChange.y)>2.5), points(log2FoldChange.y, -log10(pvalue.y), pch=19, col="red", cex = 0.75))
with(subset(res[rownames(hits),], pvalue.y<0.05 & abs(log2FoldChange.y)>0.51), points(log2FoldChange.y, -log10(pvalue.y), pch=1, col=scales::alpha("blue", 0.5), cex=0.95))
# Label points with the textxy function from the calibrate plot
with(subset(res[rownames(hits),], (pvalue.y<0.05 & abs(log2FoldChange.y)>0.51)) , textxy(log2FoldChange.y+(0.1*log2FoldChange.y), -log10(pvalue.y)+0.2*(pvalue.x), labs=labels, offset=0.5, m=c(-1.16,2), cex=0.4))
abline(h = 1.3, v = 2.5, col = "gray60", lty=3)
abline(h = 3.3, v = -2.5, col = "gray60", lty=3)
legend(1.6, 13, legend=c(expression(paste("p"["adj"]," < 0.05 & ","log"[2],"FC < |3|")), expression(paste('log'[2],'FC', ' > |2.5|')),"common hits", "NS"), col=c("red", "orange","blue","black"), pch=c(19,19,1,19), pt.cex=0.5, cex=0.5)
dev.off()


############# 6 Boxplots ######

resNvB <- results(dseqNvBvS, tidy=TRUE, name="Strain_BALB.c_vs_NOD", alpha=0.05) %>%
   arrange(padj, pvalue) %>%
   tbl_df()
resNvB

resNvF <- results(dseqNvBvS, tidy=TRUE, name="Strain_NODF_vs_NOD", alpha=0.05) %>%
   arrange(padj, pvalue) %>%
   tbl_df()
resNvF

res <- full_join(resNvF, resNvB,  by='row')

tcounts1 <- t(log2((counts(dseqNvBvS, normalized=TRUE, replaced=FALSE)+.5))) %>% 
   merge(colData(dseqNvBvS), ., by="row.names") %>%
   gather(gene, expression, (ncol(.)-length(resNvB$row)+1):ncol(.))

tcounts1 %>% 
   select(Row.names, Strain, gene, expression) %>% 
   head %>% 
   knitr::kable()


#colnames(tcounts1)[2] <- "Strain"

#tcounts <- tcounts1
#tcounts <- as.data.frame(tcounts[order(tcounts$gene),])
#tcounts$Sample <- factor(tcounts$Row.names, levels = c("F1","F2","F3","B1","B2","B3","B4","B5", "N1","N2","N3", "N4","N5"))

tcounts1$Strain <- factor(tcounts1$Strain, levels = c("NOD F","BALB/c",  "NOD M"))
tcounts1$gene <- as.factor(tcounts1$gene)
tcounts1 <- as.data.frame(tcounts1[order(tcounts1$gene),])
tcounts1$Strain <- factor(tcounts1$Strain, levels = c("NOD F", "BALB/c", "NOD M"))
tcounts1[which(tcounts1$gene=="mmu-miR-146a-5p"),]
which(levels(tcounts1$gene)=="mmu-miR-146a-5p")

chart_design <- theme(
   plot.title = element_text(color = "Black", size = 24, face = "bold", margin = margin(b=25), hjust=0.4),
   axis.text.x = element_text(size=19),
   axis.text.y = element_text(size=19),
   axis.title.x = element_text(size=18, margin = margin(t=20), colour="red2"),
   legend.text = element_blank(),
   legend.title = element_blank(),
   axis.title.y = element_text(size=20, margin = margin(r = 20)),
   strip.text.x = element_text(size =20, margin = margin(b=20), face='bold', hjust=0.4),
   strip.background = element_blank(), 
   strip.placement = "outside")

hits_final <- c("mmu-miR-322-5p","mmu-miR-146b-5p","mmu-miR-146a-5p","mmu-miR-322-3p","mmu-miR-147-3p","mmu-miR-421-3p","mmu-miR-542-3p","mmu-miR-503-5p","mmu-miR-107-3p","mmu-miR-150-5p","mmu-miR-181a-5p","mmu-miR-181b-5p","mmu-miR-3572-3p","mmu-miR-203-3p", "mmu-miR-335-3p", "mmu-miR-3572-5p", "mmu-miR-3963", "mmu-miR-3076-3p", "mmu-miR-3473a", "mmu-miR-24-2-5p", "mmu-miR-6481", "mmu-miR-1249-3p", "mmu-miR-486a-5p", "mmu-miR-486b-5p", "mmu-miR-345-5p", "mmu-miR-29b-2-5p", "mmu-miR-451a", "mmu-miR-155-5p", "mmu-miR-375-3p")

par(mfrow=c(2,5))
for (i in 1:20) {
   filename <- paste(hits_final[i], ".finalz.tiff", sep="")
   p <- ggplot(tcounts1[which(tcounts1$gene==hits_final[i]),], aes(Strain, y=expression, color=Strain)) + 
      geom_boxplot(outlier.shape = NA, width = 0.4, coef=1.8, show.legend = F, size=1.5) +
      geom_jitter(aes(fill=Strain), alpha =0.7, size=2.8, show.legend = F, width=0.2) +
      stat_summary(fun=mean, geom="point", size=2.5, color="black", alpha = 1) + 
      scale_colour_jco() +
      theme_classic() +
      ylab("log Normalized Counts")  + labs(title=paste(hits_final[i]), hjust=0.5) +
      chart_design
   
   tiff(filename, units="in", width=4.6, height=4, res=300)
   print(p)
   dev.off()
}

############## 7 NVBVF Venn Diagram ##########

NODM_ <- read.csv('NOD_norm_counts.csv', header=T, row.names = 1)
NODM_ <- NODM_[rowSums(NODM_>0)>2,]
NODM_ <- rownames(NODM_)
NODM_ <- as.list(NODM_)

BALB_ <- read.csv('Balb_norm_counts.csv',header=T,row.names = 1)
BALB_ <- BALB_[rowSums(BALB_>0)>2,]
BALB_ <- rownames(BALB_)
BALB_ <- as.list(BALB_)

NODF_ <- read.csv('F_tearNormcounts.csv',header=T,row.names = 1)
NODF_ <- NODF_[rowSums(NODF_>0)>1,]
NODF_ <- rownames(NODF_)
NODF_ <- as.list(NODF_)

library(VennDiagram)
try(library("devtools"), install.packages("devtools")) # used to source functions from the internet
library("devtools")
source_url("http://raw.github.com/nielshanson/mp_tutorial/master/downstream_analysis_r/code/venn_diagram2.r")
source_url("http://raw.github.com/nielshanson/mp_tutorial/master/downstream_analysis_r/code/venn_diagram3.r")
source_url("http://raw.github.com/nielshanson/mp_tutorial/master/downstream_analysis_r/code/venn_diagram4.r")
tiff("NvBvFatleast1.jpeg", units="in", width=6, height=5.3, res=300)
venn_diagram3(NODF_, BALB_, NODM_, name_a="NOD F",name_b ="BALB/c",  name_c = "NOD M") + scale_color_jco()
dev.off()

cpm <- count_table/colSums(count_table) * 1000000
cpm <- cpm[rowSums(cpm) > 0,]
nodm_cpm <- cpm[rowSums(cpm[,6:10]>1)>2,]
nodf_cpm <- cpm[rowSums(cpm[,11:13]>1)>1,]
balb_cpm <- cpm[rowSums(cpm[,1:5]>1)>2,]

nodm_cpm <- rownames(nodm_cpm)
nodm_cpm <- as.list(nodm_cpm)
nodf_cpm <- rownames(nodf_cpm)
nodf_cpm <- as.list(nodf_cpm)
balb_cpm <- rownames(balb_cpm)
balb_cpm <- as.list(balb_cpm)
tiff("NvBvFatleast1.jpeg", units="in", width=6, height=5.3, res=300)
venn_diagram3(nodf_cpm, balb_cpm, nodm_cpm, name_a="NOD F",name_b ="BALB/c",  name_c = "NOD M") + scale_color_jco()
dev.off()
```
